# The hippocampus as a sorter and reverberatory integrator of sensory inputs

**DOI:** 10.1101/2022.01.10.475731

**Authors:** Masanori Nomoto, Emi Murayama, Shuntaro Ohno, Reiko Okubo-Suzuki, Shin-ichi Muramatsu, Kaoru Inokuchi

## Abstract

In entorhinal-hippocampal networks, the trisynaptic pathway, including the CA3 recurrent circuit, processes episodes of context and space^1-3^. Recurrent connectivity can generate reverberatory activity^4-6^, an intrinsic activity pattern of neurons that occurs after sensory inputs have ceased. However, the role of reverberatory activity in memory encoding remains incompletely understood. Here we demonstrate that in mice, synchrony between conditioned stimulus (CS) and unconditioned stimulus (US)-responsible cells occurs during the reverberatory phase, lasting for approximately 15 s, but not during CS and US inputs, in the CA1 and the reverberation is crucial for the linking of CS and US in the encoding of delay-type cued-fear memory. Retrieval-responsive cells developed primarily during the reverberatory phase. Mutant mice lacking N-methyl-D-aspartate receptors (NRs) in CA3 showed a cued-fear memory impairment and a decrease in synchronized reverberatory activities between CS- and US-responsive CA1 cells. Optogenetic CA3 silencing at the reverberatory phase during learning impaired cued-fear memory. Our findings suggest that reverberation recruits future retrieval-responsive cells via synchrony between CS- and US-responsive cells. The hippocampus uses reverberatory activity to link CS and US inputs, and avoid crosstalk during sensory inputs.

---

The hippocampus, a centre of multimodal convergence, is crucial for learning and memory of associative episodes^7^. The hippocampus is primarily comprised of four subfields: the dentate gyrus (DG), CA3, CA2, and CA1. There are two parallel pathways, the trisynaptic pathway and the monosynaptic pathway. In the trisynaptic pathway, which is important for one-trial contextual learning, information flows from the entorhinal cortex (EC) to the DG, to CA3, to CA1, and finally to the EC^1-3^. In the monosynaptic pathway, which is important for temporal association, information flows from the EC to CA1 to the EC^8^. Among these pathways, the CA3 has a unique system, a recurrent circuit forming extensive interconnections within CA3 cells^9^. The CA3 recurrent circuit is crucial for pattern completion^10^, an ability of recall from partial cue. In addition, theoretical models have suggested that the CA3 recurrent circuit implemented with NR function has a potential to generate reverberatory neuronal activities without input from external stimuli, and acts as an associator of temporally separated episodes by filling a temporal gap between discontinuous events^11-15^. However, experimental studies have indicated that the CA3 recurrent circuit of the trisynaptic pathway is not required for trace-type associative memory formation, which requires the ability to form temporal association between events^8, 16, 17^. Therefore, the roles in memory association of potential reverberatory activities in the hippocampus remain to be elucidated. The entorhinal-hippocampal network contains a microcircuit that restricts incorporation of US input as a part of CS representation during sensory inputs^18^. We hypothesized that the hippocampal network is programmed to process sensory information after termination of sensory inputs and reverberatory activities, which could represent the CS and the US, serve as an integrator to link neutral and aversive stimuli. We sought to determine whether and how reverberatory activities contribute to the CS-US association.

We subjected mutant mice that specifically lack NRs in CA3 (CA3-NR1 KO mice), and thus are deficient in NR current and synaptic plasticity at the recurrent CA3 synapses^10^, to a delayed-type light-cued-fear conditioning (LFC) task^19^ (Fig. 1a, b). CA3-NR1 KO mice and littermate controls were both subjected to pre-contextual habituation followed by training sessions to form an association between a light-cue CS and a footshock US with ten pairings. In the following test, CA3-NR1 KO mice exhibited impairment in long-term, but not short-term, cued-fear memory recall relative to littermate controls (Fig. 1c, d). By contrast, CA3-NR1 KO mice did not exhibit impairments in long-term contextual fear memory in the LFC task (Fig. 1e) or in an alternative contextual fear memory task, which consisted of context pre-exposure and immediate footshock sessions^20, 21^ (Extended Data Fig. 1a, b). Our results testing contextual fear memories in CA3-NR1 KO mice confirmed previous findings, which suggested that CA3 NRs are important for novel contextual representation but not familiarised context representation^1-3^. Furthermore, long-term memory recall of an auditory-cued-fear conditioning (AFC) task was impaired in CA3-NR1 KO mice relative to controls (Fig. 1f, g), indicating that CA3 NRs are important for long-term cued-fear memory.

**Figure 1.**
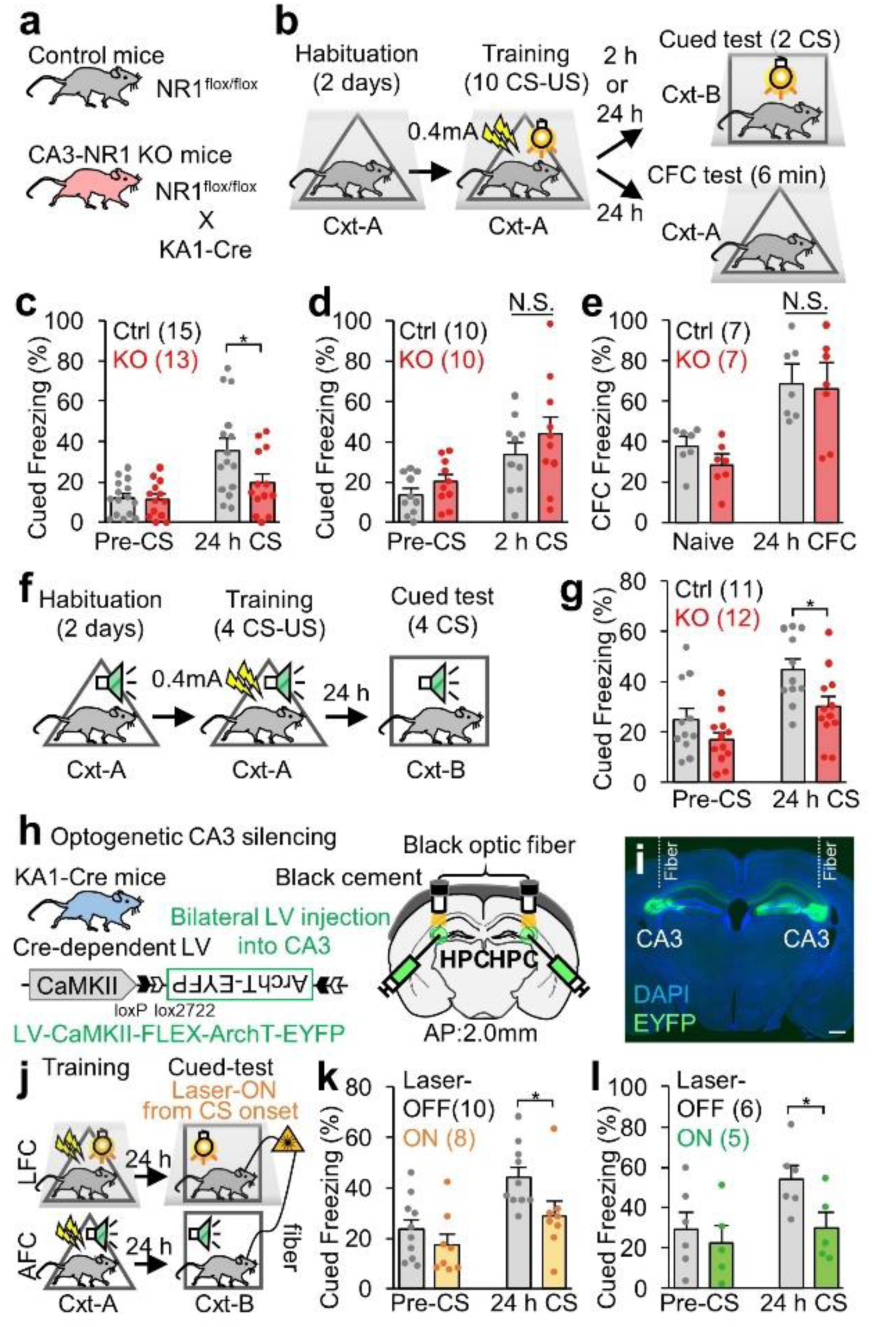
CA3 NRs and CA3 activity are important for cued-fear memory. **a,** Animals used in this study. **b,** Experimental design for light-cued-fear conditioning (LFC) task. **c, d,** Cued freezing levels during (**c**) 24 h long-term, and (**d**) 2 h short-term, memory tests. **e,** Contextual freezing levels during a 24 h long-term memory test in the LFC task. **f,** Experimental design for auditory-cued-fear conditioning (AFC) task. **g,** Cued freezing levels during the 24 h long-term test in AFC task. **h,** Animal and virus vector used for optogenetic CA3 silencing. **i,** Coronal section of the hippocampus with EYFP-expressing cells. Scale bar, 500 μm. **j,** Experimental design for optogenetic experiment. **k, l** Cued freezing levels during the 24 h long-term memory test in the (**k**) LFC, and (**l**) AFC tasks. **c-e, g, k, l,** *P* values determined using an unpaired two-tailed *t* test (\**P* < 0.05). Graphs represent the mean ± SEM, and circles within the graphs represent individual animals. Numbers in parentheses denote the number of mice in each group used for the study. Lightning bolt, footshock; Light bulb, light CS; Cxt, context; Speaker, tone CS; HPC, hippocampus; AP, anterior-posterior; N.S., not significant.

Lentivirus (LV) encoding calcium/calmodulin-dependent protein kinase II (CaMKII)-FLEX-eArch3.0-EYFP was bilaterally injected into the hippocampal CA3 of KA1::Cre heterozygous transgenic mice to specifically label and silence CA3 excitatory cells (Fig. 1h, i). One day after the training session, mice were subjected to test sessions during which a continuous laser (589 nm) was bilaterally delivered to the CA3, starting at the onset of the first CS in the LFC or AFC tasks (Fig. 1j). Mice with precise optical CA3 silencing exhibited impaired long-term cued-fear memory recall in both LFC and AFC tasks relative to the control (laser-OFF) group (Fig. 1k, l). These results indicated that NRs and neuronal activities in the CA3 are important for cued-fear memory, which is consistent with previous reports^22-24^ indicating involvement of the hippocampus in cued-fear memory.

To investigate how the hippocampus processes CS and US information, and whether reverberatory activity emerges after termination of sensory stimuli in the cued- fear conditioning task, we monitored *in vivo* transient calcium (Ca^2+^) dynamics in hippocampal cells. CA3-NR1 KO mice and littermate controls were injected with an Adeno-Associated Virus (AAV) encoding CaMKII-G-CaMP7 and implanted with a micro-gradient index (GRIN) lens targeting the right CA1 (Fig. 2a). The same CA1 cells were tracked across LFC task sessions, including a 30 min rest session after the training, using an automated sorting system to extract the Ca^2+^ activity of each neuron, which was then normalized using z-scores (Extended Data Fig. 2).

**Figure 2.**
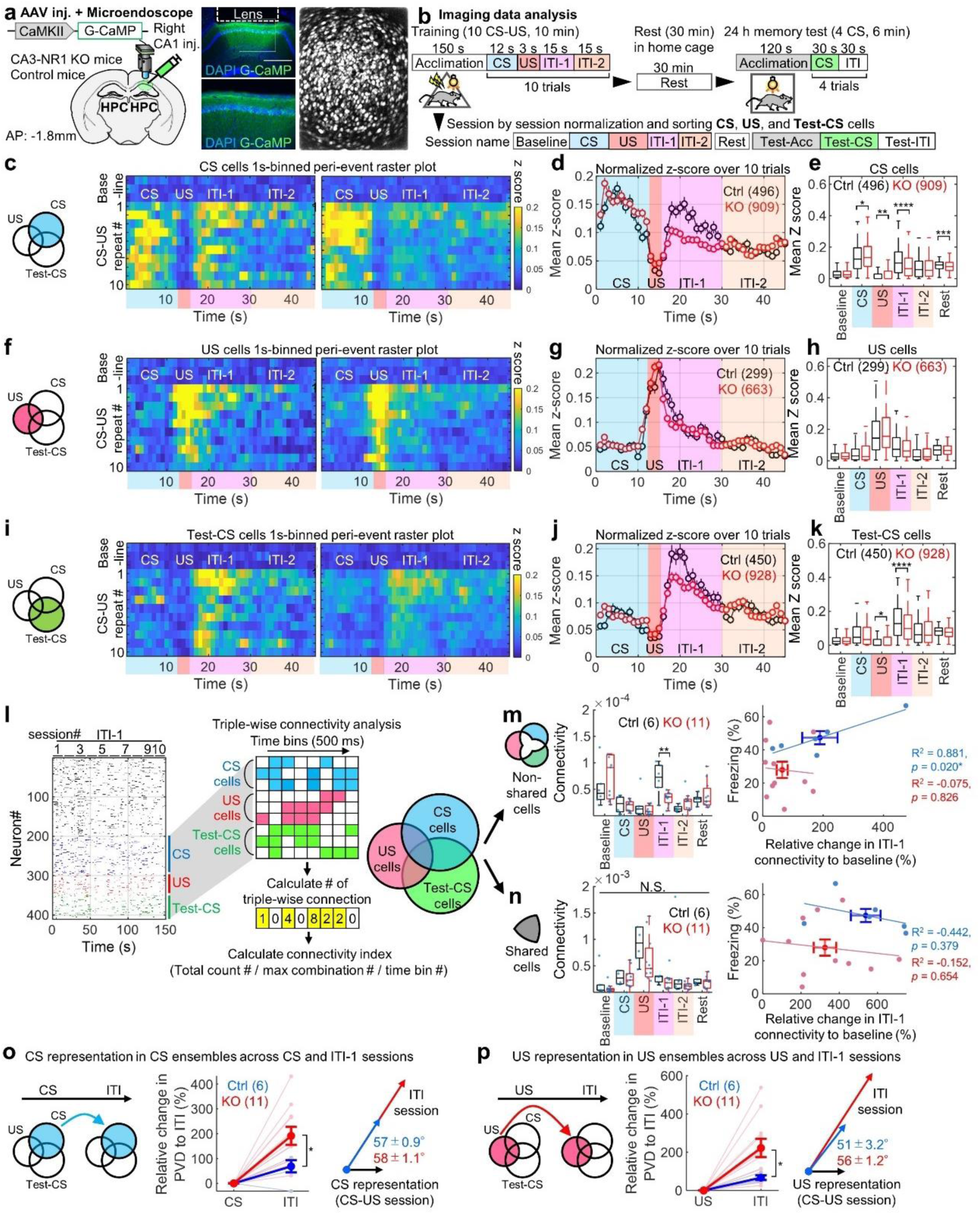
CA3 NRs are involved in reverberatory and synchronized activity, but not sensory propagation, in CS- and Test-CS-responsive ensembles. **a,** Left, experimental design. Right, coronal section of the hippocampus with G-CaMP expression, GRIN lens implantation, and stacked dF/F image acquired using a microendoscope over entire recording sessions of hippocampal imaging. Scale bar, 500μm. **b,** Imaging data analysis scheme. In each cell, Ca^2+^ data is classified into nine sessions, and the calculated mean z-score is considered to represent responsiveness and sorted into CS-, US-, and Test-CS-responsive subpopulations. **c,** Venn diagrams representing CS ensembles. Peri-event raster plots during the training session in CS- responsive subpopulations of wild-type littermate (left) control and (right) KO mice. Each short vertical tick represents a 1 s change of mean z-score across baseline and ten CS-US pairings. Ca^2+^ activities were aligned at the time that CS-US stimuli were delivered. The color code represents mean z-score. **d,** Averaged z-score plots over ten CS-US pairings in CS-responsive subpopulations. **e,** Box plots comparing mean z- scores between genotypes in each session. **f,** Venn diagrams representing US ensembles. Peri-event raster plots during the training session in US-responsive subpopulations of wild-type littermate (left) control and (right) KO mice. Each short vertical tick represents a 1 s change of mean z-score across baseline and ten CS-US pairings. Ca^2+^ activities were aligned at the time that CS-US stimuli were delivered. The color code represents mean z-score. **g,** Averaged z-score plots over ten CS-US pairings in US-responsive subpopulations. **h,** Box plots comparing mean z-scores between genotypes in each session. **i,** Venn diagrams representing Test-CS ensembles. Peri-event raster plots during the training session in Test-CS-responsive subpopulations of wild-type littermate (left) control and (right) KO mice. Each short vertical tick represents a 1 s change of mean z-score across baseline and ten CS-US pairings. Ca^2+^ activities were aligned at the time that CS-US stimuli were delivered. The color code represents mean z-score. **j,** Averaged z-score plots over ten CS-US pairings in Test-CS-responsive subpopulations. **k,** Box plots comparing mean z-scores between genotypes in each session. **l,** Left, representative binarized raster plots of Ca^2+^ activity across ten ITI-1 sessions in control animals. Right, magnified raster plots focusing on CS-, US-, and Test-CS-responsive subpopulations and scheme for connectivity analysis. This analysis calculates connectivity by normalizing the number of synchronized connections every 500 ms among the three subpopulations in each session. **m, n,** Box plots comparing mean connectivity between genotypes in each session. **o, p,** Mahalanobis PVD and rotation between CS and ITI-1 sessions in the (o) CS-responsive ensemble and between US and ITI-1 sessions in the (p) US-responsive ensemble. Numbers in parentheses denote the number of (**d, e, g, h, j, k**) cells or (**m, n, o, p**) mice in each group used for the studies. *P* values were determined using a (**e, h, k**) Wilcoxon rank sum test, (**m, n, o, p**) Unpaired *t* test, or (**m, n**) Pearson correlation (\**P* < 0.05, \*\**P* < 0.01, \*\*\**P* < 0.001, \*\*\*\**P* < 0.0001). Box plots represent median, first, and third quantiles, and minimum and maximum values. Graphs represent means ± SEM.

By calculating and comparing mean z-scores as indicators of the responsiveness for each session (please refer to Methods), we sorted training CS (CS)-, training US (US)-, and long-term memory-test CS (Test-CS)-responsive subpopulations of cells that exhibited 2-fold higher responses to stimuli than in the baseline session on the training day before the corresponding CS or US (Fig 2b, c, f, i). We did not detect structural differences of these CA1 subpopulations between CA3-NR1 KO mice and littermate controls (Extended Data Fig. 3a-c). About half of the Test-CS-responsive cells had newly emerged and were not CS-responsive cells, indicating that this cell subpopulation that responded to the CS changed from the training session to the test session (Extended Data Fig. 3e). Notably, US input immediately and completely shut down the activity of CS-responsive and Test-CS-responsive cells (Fig 2b-e, i-k). The Ca^2+^ activities of the Test-CS-responsive cells, but not the CS-responsive or US-responsive cells, during CS presentation in the test session were higher than those in the corresponding acclimation (Test-Acc) and inter-trial interval (Test-ITI) sessions (Extended Data Fig. 3d). CS- responsive cells in the CA3-NR1 KO mice exhibited dramatically decreased Ca^2+^ activity relative to the control during the ITI, especially during the first 15 s after the CS-US presentation (ITI-1), and also during the 30 min rest session (Fig. 2c-e, Extended Data Fig. 4, Supplementary Movie 1). The activities of US-responsive cells were comparable between genotypes throughout all sessions (Fig. 2f-h). Ca^2+^ activity of the Test-CS-responsive cells emerged after CS-US presentation (Fig. 2i-k). CS- responsive and Test-CS-responsive cells in CA3-NR1 KO mice exhibited less activity during the ITI-1 than the control. Together, these studies indicated that in the CA1, ablation of CA3 NRs decreases reverberatory activities in current (CS) and future (Test-CS) CS-responsive cell ensembles, which emerge after termination of sensory stimuli, without affecting the ensemble structure.

**Figure 3.**
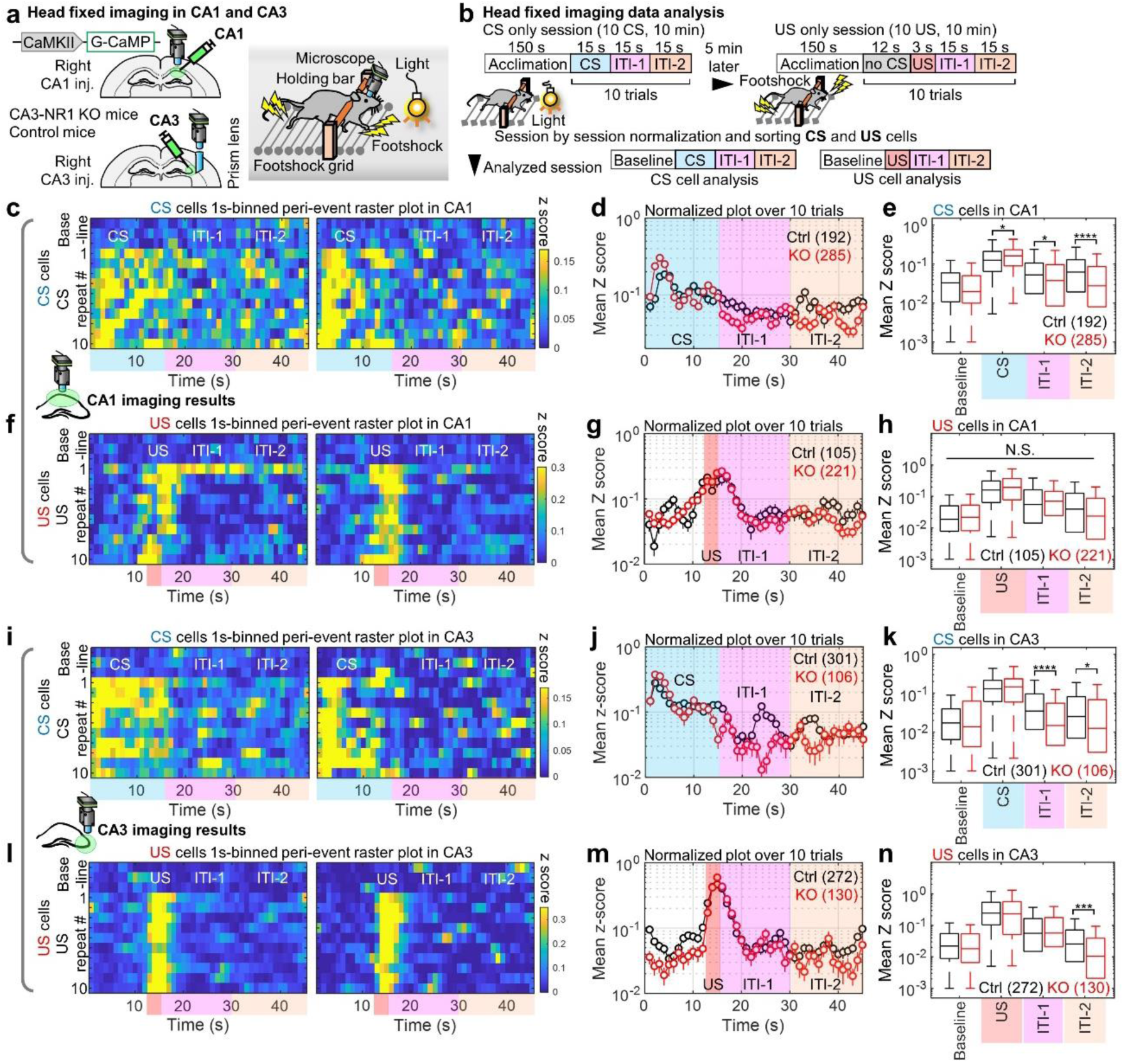
CA3 NRs-dependent reverberation by single stimuli in the hippocampal network under head-fixed conditions. **a,** Left, experimental design. Right, schema for head-fixed imaging. The same footshock grid and light bulb were used in free-moving and head-fixed imaging experiments. **b,** Imaging data analysis scheme. In each cell, Ca^2+^ data are classified into four sessions, and the calculated mean z-score, considered to represent responsiveness, is sorted into CS- and US-responsive subpopulations. **c,** Peri-event raster plots during single CS presentation session in CA1 subpopulations of (left) control and (right) KO mice. Each short vertical tick represents a 1 s change of mean z-score across baseline, and CS presentations. Ca^2+^ activities are aligned at the time at which stimuli were delivered. The color code indicates mean z-score. **d,** Averaged z-score plots over ten CS presentations in CA1 CS-responsive subpopulations. **e,** Box plots comparing mean z-scores between genotypes in each session. Numbers in parentheses denote the number of cells in each group used for the study. **f,** Peri-event raster plots during single US presentation session in CA1 subpopulations of (left) control and (right) KO mice. Each short vertical tick represents a 1 s change of mean z-score across baseline, and US presentations. Ca^2+^ activities are aligned at the time at which stimuli were delivered. The color code indicates mean z-score. **g,** Averaged z-score plots over ten US presentations in CA1 US-responsive subpopulations. **h,** Box plots comparing mean z-scores between genotypes in each session. Numbers in parentheses denote the number of cells in each group used for the study. **i,** Peri-event raster plots during single CS presentation session in CA3 subpopulations of (left) control and (right) KO mice. Each short vertical tick represents a 1 s change of mean z-score across baseline, and CS presentations. Ca^2+^ activities are aligned at the time at which stimuli were delivered. The color code indicates mean z-score. **j,** Averaged z-score plots over ten CS presentations in CA3 CS-responsive subpopulations. **k,** Box plots comparing mean z-scores between genotypes in each session. Numbers in parentheses denote the number of cells in each group used for the study. **l,** Peri-event raster plots during single US presentation session in CA3 subpopulations of (left) control and (right) KO mice. Each short vertical tick represents a 1 s change of mean z-score across baseline, and US presentations. Ca^2+^ activities are aligned at the time at which stimuli were delivered. The color code indicates mean z-score. **m,** Averaged z-score plots over ten US presentations in CA3 US-responsive subpopulations. **n,** Box plots comparing mean z-scores between genotypes in each session. Numbers in parentheses denote the number of cells in each group used for the study. Data were acquired from Ctrl (n = 2 mice) and KO (n = 2 mice) groups for CA1 imaging, and from Ctrl (n = 4) and KO (n = 2 mice) groups for CA3 imaging. *P* values were determined using (e, h, k, n) a Wilcoxon rank sum test (\**P* < 0.05, \*\**P* < 0.01, \*\*\**P* < 0.001, \*\*\*\**P* < 0.0001). Box plots represent the median, first, and third quantiles, and minimum and maximum values. Graphs represent means ± SEM.

**Figure 4.**
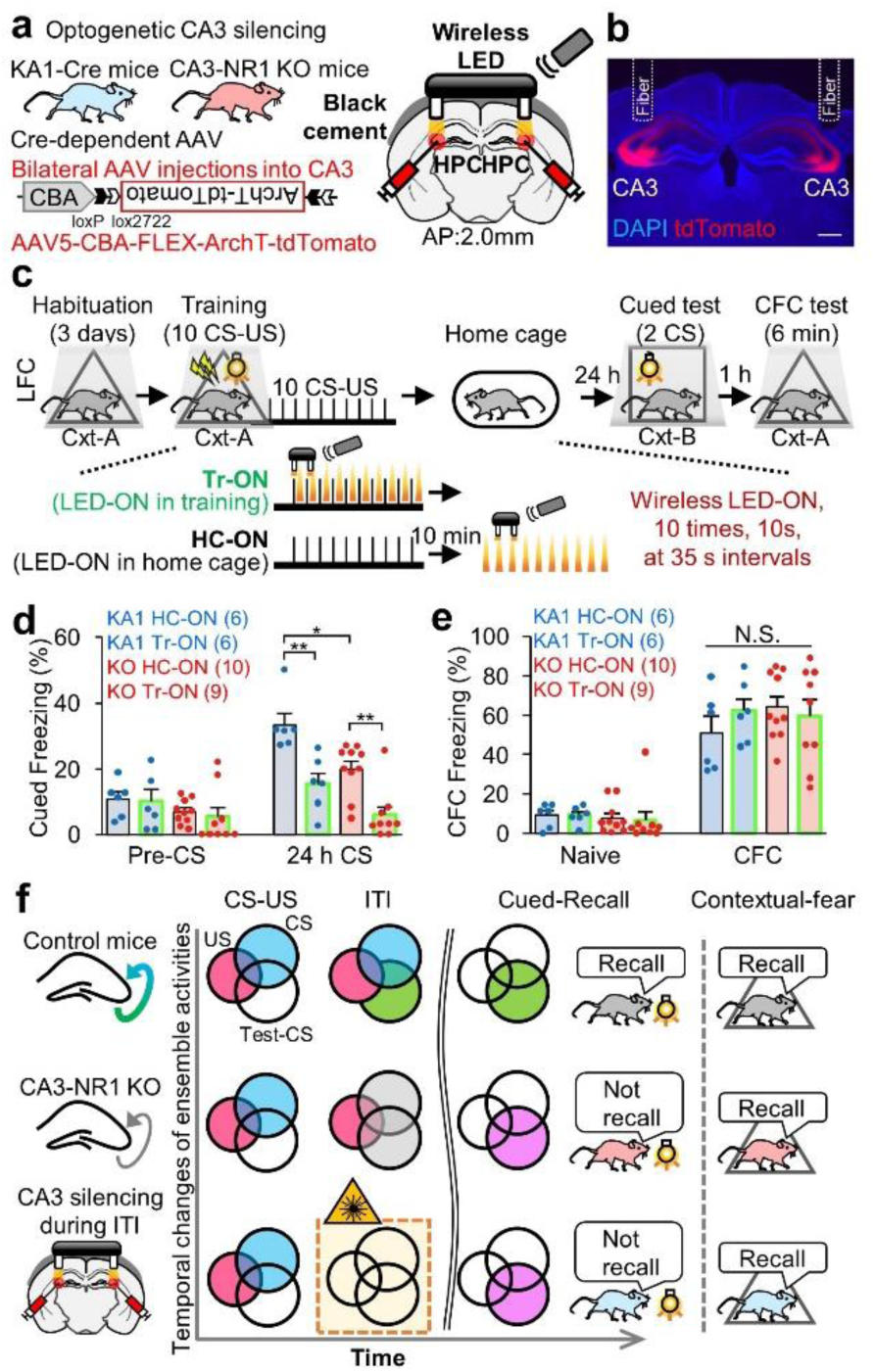
CA3-CA1 pathway activity after termination of sensory stimuli is crucial for cued-fear memory encoding but not for contextual fear memory. **a,** Experimental design. **b,** Coronal section of the hippocampus with tdTomato expression and fiber implantation targeting CA3. Scale bar, 500 μm. **c,** Scheme for optogenetic manipulation. After the habituation session, mice were subjected to ten CS-US pairings in a training session. The CA3-CA1 pathway was silenced ten times for 10 s at 35 s intervals, during either the ITI phase following CS-US presentation (Tr-ON) or during resting in the home cage 10 min after the training session (HC-ON). On the next day, mice were tested for cued freezing and contextual freezing. **d,** Cued freezing levels during the 24 h long-term memory test. **e,** Contextual freezing levels during the 24 h long-term memory test. *P* values were determined using a two-way analysis of variance (ANOVA) with the Tukey–Kramer test (\**P* < 0.05, \*\**P* < 0.01). ANOVA of cued freezing level: genotype. Graphs represent means ± SEM, with circles indicating individual animals. **f,** Summarized scheme of imaging and behavioral results in the study. Venn diagrams of temporal changes in ensemble activities corresponding to CS, US, and Test-CS. Filled circles with color indicate the activated ensemble in each behavioral session of experimental groups. The CA3-NR1 KO group exhibited less activity in the CS- and Test-CS-responsive ensembles during ITI and failed cued-fear memory recalls. Numbers in parentheses denote the number of mice in each group used for the study. Lightning bolt, footshock; Light bulb, light CS; Cxt, context; HPC, hippocampus; AP, anterior-posterior; N.S., not significant.

The postulated advantage of the reverberatory activity is to prolong the time window that allows temporal coordination among cell ensembles to integrate stimuli, leading to associative memory formation^4-6^. To determine if synchronized activity among cell ensembles increases during the ITI-1, we counted the number of pairwise synchronized Ca^2+^ activities within 500 ms (please refer to Methods) (Extended Data Fig. 5). CA3-NR1 KO mice exhibited significantly lower pairwise connectivity between newly generated Test-CS-responsive cells (Test-CS-specific cells) and (CS US)- responsive cells during the ITI-1 than the control group. Furthermore, triple connectivity between CS-, US-, and Test-CS-specific cells in CA3-NR1 KO mice was significantly lower than that of littermate controls only during the ITI-1 (Fig. 2l, m). Notably, the control group exhibited a positive correlation between the relative degree of animals’ freezing on stimulus and the degree of ITI-1 connectivity. These characteristic features of connectivity were not observed in shared cells (Fig. 2n).

To further assess the representation similarity between CS and US cell ensembles across CS, US, and ITI-1 sessions, we calculated the Mahalanobis population vector distance (PVD)^25^ following principal component analysis (PCA)-based dimension reduction^26^ and rotation of multidimensional population vectors (one dimension per cell)^25^ (Fig. 2o, p). CS or US ensembles in CA3-NR1 KO mice and littermate controls exhibited comparable rotation from CS or US to ITI (Fig. 2o, p). Both CS and US ensemble representations were more stable in littermate control mice than in CA3-NR1 KO mice across sessions, as demonstrated by small PVD changes from the stimulus to the ITI session. This suggests that reverberatory activities repeat CS and US representations. By contrast, ensemble activities exhibited increased variation across sessions in CA3-NR1 KO mice.

The hippocampus processes multimodal information and contains a wide variety of cell types, such as place and head-direction cells^27^, which could contribute to the observed reverberatory activity. Thus, we determined if the sensory stimuli alone (CS or US) triggers reverberatory activity in the hippocampal network using head-fixed mice. Mice operated for imaging in the CA1 (Fig. 2a) and the CA3 (Extended Data Fig. 7a) were additionally prepared in such a manner as to allow head fixation on a head-fixed apparatus consisting of a footshock grid and a light bulb via a holding bar with dental cement (Fig. 3a). After habituation for 4 days, mice were subjected to a training session in which they were exposed to the CS or the US alone using the same exposure time and interval as in the LFC task (Fig. 3b). The training CS- and US-responsive subpopulations were sorted using the same criteria (2-fold higher responses to stimuli). The CS-responsive cells in both the CA1 and CA3 of CA3-NR1 KO mice exhibited significantly lower Ca^2+^ activity than the littermate controls during the ITI-1 and the ITI-2 (Fig. 3c-e, i-k, Extended Data Fig. 6, Supplementary movie 2). By contrast, the US-responsive cells exhibited comparable activities in both genotypes (Fig. 3f-h, l-n). These findings, combined with the findings in freely moving mice, demonstrate that during sensory input, CA3 NRs are not important for direct propagation of CS and US information into the hippocampal CA3-CA1 network, but rather are crucial for reverberation of CS, but not US, representation in this network.

We also examined CA3 dynamics during the LFC task in freely moving conditions (Extended Data Fig. 7a, b, c, f, i). There were no significant differences in the cell ensemble structure or in the Ca^2+^ activities during test sessions between CA3- NR1 KO mice and littermate controls (Extended Data Fig. 8a-d). In contrast to the CA1, US input enhanced, rather than inhibited, the Ca^2+^ activities of CS- and Test-CS- responsive cells until the early half period of the ITI-1 (1 to 5 s of 15 s in ITI-1). The CS- and Test-CS-responsive cells in CA3-NR1 KO mice exhibited significantly lower Ca^2+^ activities during the subsequent 9 s of the ITI-1 (6 to 14 s of 15 s in ITI-1) and ITI- 2 sessions (Extended Data Fig. 7c-e, i-k) than the control. Activities of US-responsive cells were comparable between genotypes (Extended Data Fig. 7f-h). Triple connectivity was comparable between CA3-NR1 KO and littermate control mice for the duration of sessions (Extended Data Fig. 7l-n). Mahalanobis PVD analysis revealed comparable similarities in CS and US ensemble representations across CS-US and ITI sessions between genotypes (Extended Data Fig. 7o, p). The CS ensemble, but not the US ensemble, exhibited modest but statistically significant rotation from the stimulus to the ITI in CA3-NR1 KO mice relative to littermate controls (Extended Data Fig. 7o, p). These hippocampal CA3 and CA1 imaging results suggested that after termination of sensory stimuli, the CA3-CA1 pathway acts as a reverberatory network of episodes in a CA3 NR-dependent manner.

Finally, we determined if CA3 reverberatory activity is crucial for the association between the CS and US. KA1::Cre/CA3-NR1-KO mice were bilaterally injected in the CA3 with an AAV encoding chicken beta actin (CBA)-FLEX-ArchT- tdTomato to specifically label CA3 cells with ArchT-tdTomato. Wireless optogenetic LED (590 nm) cannulae were implanted bilaterally into the CA3 (Fig. 4a, b). CA3 neuronal activity was optogenetically silenced either during the ITI for 10 s immediately after CS-US presentation (Tr) or during a 10 min rest period after the training session (HC) in cued-fear conditioning using the same silencing intervals (please refer to Methods) (Fig. 4c). One day after the training session, mice were subjected to a cued- fear memory test followed by a contextual fear memory test at a 1 h interval (Fig. 4d, e). Consistent with the behavioral data in Fig. 1, CA3-NR1 KO mice exhibited impaired cued-fear memory and unchanged contextual memory recall compared with KA1::Cre mice (KA1 HC-ON vs KO HC-ON) (Fig. 4d, e). Importantly, mice that received silencing at the time of the early reverberatory phase (Tr-ON) exhibited significantly decreased cued-fear recall, but similar contextual fear memory in both genotypes compared with the group silenced after training (HC-ON). Taken together, these findings suggested that CA3 reverberatory activity is crucial for cued-fear memory encoding (Fig. 4f).

We detected time-limited and CA3 NR-dependent reverberatory activities that lead to synchronized activity among cell ensembles in the CA1. The CA3 to CA1 network functions as a reverberatory and associative system of stimuli, in which the CA3 acts as a reverberator and the CA1 functions as both a reverberator and an integrator of episodes.

This prompts the question of why CS and US events must interact during the reverberatory phase. Simultaneous encoding of multiple stimuli in the brain neural network is limited by the capacity of cognition^28^. The hippocampus processes distinct valences, such as contextual and temporal episodes, with distinct cell subpopulations^29–, 32^ and circuits^1-3, 8^. The hippocampus could use reverberatory activity to avoid crosstalk during sensory inputs to store neutral and aversive information separately, and subsequently link the CS and US during the reverberatory phase. Indeed, a circuit mechanism in which CA1 activity is temporally regulated by EC input prevents crosstalk between CS and US stimuli during contextual CS and aversive US presentations^18^. Thus, the hippocampus functions as a sorter to encode CS and US independently, and subsequently as a reverberatory integrator to link CS and US.

CS-responsive cells occupied about half of Test-CS-responsive cells, that is, another half of the Test-CS cells emerged after the conditioning. The significant correlation between relative degree of animal freezing and triple connectivity of non- shared cells during ITI strongly suggests that, by synchronized activity, CS and US cells recruit and instruct newly generated Test-CS cells by synchronizing CS and US information (Extended Data Fig. 9). The hippocampus allocates CS and US events into distinct cell subpopulations (Fig. 4f). In the amygdala, the CS-responsive cell ensemble begins to respond to US stimuli across repetitive CS-US presentations and eventually represents the US to encode cued-fear memory^25^. In addition, the amygdala exhibits greater overlapping in cell populations between training and retrieval sessions than the CA1^33^. Thus, the amygdala encodes CS-US association by altering the valence representation of the cell ensemble. Our findings suggest that the hippocampus and amygdala adopt different strategies to encode distinct aspects of associative memory, episodic relation and direct linking, respectively.

Human studies reveal that in delayed conditioning in which the neutral cue (CS) is paired with the aversive stimulus (US), the hippocampus and amygdala act in parallel to associate the CS and US^34, 35^. The hippocampus is crucial for declarative association between the CS and US, while the amygdala is involved in automatic conditioned responses. In rodents, general theory indicates that the hippocampus is dispensable for delayed conditioning^36, 37^. However, when conditioning stimuli are weaker and do not trigger the amygdala as robustly, hippocampal contribution to behavior becomes apparent^22, 23, 38, 39^. We suggest that, similar to humans, the mouse hippocampus integrates the episodic relation between the CS and US, while the amygdala mediates direct associations between the CS and US (Extended Data Fig. 10).

Consistent with previous reports^1-3^, NR deficiency in the CA3 or CA3 silencing during reverberation did not impair contextual fear memories after pre-contextual habituation (Fig. 1e, Fig. 4e, Extended Data Fig. 1a, b). These findings suggest that reverberation is required for association of novel episodes but not for association with pre-existing memories.

The slow kinetics of NRs are thought to be crucial for holding evoked excitation within the recurrent network^14, 15^. Therefore, reverberatory activity is initially generated through the CA3 recurrent circuit in an NR-dependent manner. The entorhinal- hippocampal time-limited gate opens immediately after the sensory stimulus^18, 40^, which promotes propagation to the CA1 and initiates reverberation. Synchronized activities between CS- and US-responsive cells is stochastically regulated during the ITI, generating novel Test-CS-responsive cells.

## Supporting information

Summary of statistical analyses in main and extended figures.

## Methods

### Mice

Male CA3-NR1 KO mice (C57BL/6J background) and their floxed-NR1 littermate controls were used for behavioral, imaging, and optogenetic experiments. Male KA1::Cre mice were used for optogenetic experiments. CA3-NR1 KO mice were generated by crossing floxed-NR1 and KA1::Cre transgenic mice. Mice were maintained on a 12 h light-dark cycle at 24°C ± 3°C and 55% ± 5% humidity with standard laboratory diet and tap water ad libitum. All mice were aged 16–26 weeks at the time of behavioral experiments. All procedures involving the use of animals complied with the guidelines of the National Institutes of Health and were approved by the Animal Care and Use Committee of the University of Toyama, Toyama, Japan.

### Viral constructs

For the *in vivo* Ca^2+^ imaging experiment, a recombinant Adeno-Associated Virus (AAV) vector encoding AAV^9^-CaMKII::G-CaMP7 (Titer: 9.4 × 10^12^ vg/mL) after 40- fold dilution with phosphate buffered saline (PBS) (T900; Takara Bio, Inc., Japan) was used^41^. For optogenetic silencing during the training session, AAV encoding AAV5- CBA-FLEX-ArchT-tdTomato (Titer: 1.3 × 10^13^ GC/mL) (#28305; Addgene, USA) after 10-fold dilution with PBS was used. For optogenetic silencing during the test session, lentivirus (LV) encoding CaMKII-FLEX-eArch3.0-EYFP (Titer: 5 × 10^9^ IU mL^−1^) without dilution was used. The LV was prepared as described previously^29^, according to the protocol developed by K. Deisseroth.

### Stereotaxic surgery for optogenetic and imaging studies

Stereotaxic surgery and optic fiber placement were conducted as described previously^29^. Prior to surgery, mice were anesthetized with intraperitoneal injection of a three-drug combination: 0.75 mg/kg medetomidine (Domitor; Nippon Zenyaku Kogyo Co., Ltd., Japan); 4.0 mg/kg midazolam (Fuji Pharma Co., Ltd., Japan); and 5.0 mg/kg butorphanol (Vetorphale; Meiji Seika Pharma Co., Ltd., Japan). After surgery, an intramuscular injection of 1.5 mg/kg atipamezole (Antisedan; Nippon Zenyaku Kogyo), a medetomidine antagonist, was administered to reverse sedation. Mice were placed on a stereotaxic apparatus (Narishige, Japan), and subsequently bilaterally injected with LV or AAV solution into the dorsal hippocampal CA3 (from bregma: +2.0 mm anteroposterior [AP], ±2.2 mm mediolateral [ML]; from dura: +1.8 mm dorsoventral [DV]). All virus injections were conducted using a 10 μL Hamilton syringe (80030; Hamilton, USA) fitted with a mineral oil-filled glass needle and wired to an automated motorized microinjector IMS-20 (Narishige). The glass injection tip was maintained before and after injection at the target coordinates for 5 min.

For the wired optogenetic experiment, mice were bilaterally injected with 500 nL LV solution at 100 nL min^−1^ into the CA3, and bilaterally implanted with guide cannulas targeting the CA3 (from bregma: -2.0 mm AP, ±2.2 mm ML; from dura: +1.3 mm DV; C313GS-5/SPC, 22-gauge; Plastics One, USA). Dummy cannulas (C313IDCS-5/SPC, zero projection, Plastics One) were then inserted into guide cannulas to protect these guide cannula tubes from dust.

For the wireless optogenetic experiment, a wireless optogenetics system, Teleopto (Bio Research Center, Japan), was used^42^. Mice were bilaterally injected with 500 nL AAV solution at 100 nL min^−1^ into the CA3 and implanted with a dual-LED cannula (fiber diameter, 500 µm; fiber length, 3.3 mm; bilateral, 590 nm, 10 mW) targeting the CA3 (from bregma: -2.0 mm AP, ±2.2 mm ML; from dura: +1.2 mm DV). Micro-screws were fixed near the bregma and lambda, and guide cannulas were fixed in position using dental cement (Provinice; Shofu, Inc., Japan) mixed with 5% carbon powder (484164; Sigma, USA). Mice were allowed to recover from surgery for 4 weeks in their home cages before behavioral experiments were initiated.

For the Ca^2+^imaging experiment, surgery was conducted as described previously with modifications^41^. Mice were unilaterally injected with 500 nL AAV9-CaMKII::G- CaMP7 at 100 nL min^−1^ into the right hippocampal CA1 (from bregma: -2.0 mm AP, +1.4 mm ML, +1.4 mm DV) or CA3 (from bregma: -2.0 mm AP, +2.2 mm ML; from dura: +1.8 mm DV). After 1 week of recovery from AAV injection surgery, anesthetized mice were placed back onto a stereotactic apparatus to implant a gradient index (GRIN) lens into CA1 or CA3. A craniotomy (CA1, approximately 1.8 mm in diameter; CA3, approximately a 2.0 × 2.0 mm square) was performed centered over the injection site, and the neocortex and corpus callosum above the alveus overlying the dorsal hippocampal CA1 or CA3 were aspirated under constant irrigation with saline using a 26-gauge flat-blunted needle tip. Saline was applied to control bleeding. A cylindrical GRIN lens (diameter, 1.0 mm; length, 4 mm; Inscopix, USA) and prism GRIN lens (diameter, 1.0 mm; length, 4 mm with prism lens; Inscopix) were attached to the alveus and additionally squeezed 10–30 μm using handmade forceps attached to a manipulator (Narishige) for CA1 and CA3 imaging. Emulsified low-temperature bone wax was applied to seal the gaps between the GRIN lenses and the skull, and the lens was then anchored in place using dental cement mixed with 5% carbon powder (464164, Sigma) as described above. After the surgery, Ringer’s solution (0.5 mL/mouse, i.p.; Otsuka, Japan) was injected, and atipamezole was administered as described above. Mice were maintained in individual cages after surgery. Three weeks after GRIN lens implantation, mice were anesthetized and placed back onto the stereotaxic apparatus to set a baseplate (Inscopix). A Gripper (Inscopix) holding a baseplate attached to a miniature microscope (nVista 3, Inscopix) was lowered over the implanted GRIN lens until visualization of clear vasculature was possible, indicating the optimum focal plane. Carbon-containing dental cement was then applied to fix the baseplate in position and preserve the optimal focal plane. Mice recovered from surgery in their home cages at least for 1 week before beginning behavioral imaging experiments.

For CA1 and CA3 head-fixed imaging experiments, mice that had undergone the surgery until the step of baseplate setting were anesthetized and placed back onto a stereotaxic apparatus, and mice were removably attached to a holding bar with a dental cement. The cut tips of a PCR tube held with ear bar for stereotaxic (Narishige) bilaterally were moved closer to the face between the eye and ear, and then the tips were fixed with dental cement, enabling the stereotaxic device to hold the mouse head on the apparatus.

### Behavioral analysis

All mice were numbered and randomly assigned to each experimental group before the experiments, with the exception of an imaging experiment. All behavioral experiments were performed and analyzed by an investigator blinded to experimental conditions with the exception of imaging experiment. For all behavioral procedures, animals in their home cages were moved on a rack to a resting room next to the behavioral testing room and left undisturbed for at least 30 min before each behavioral experiment. All behavioral chambers were cleaned after each behavioral session. After all the optogenetic experiments were completed, the injection sites were histologically verified. Data were excluded from behavior analyses if the animals exhibited abnormal behavior after surgery, the target area was missed, or the bilateral expression of the virus was inadequate. All behavioral sessions were conducted using a video tracking system (Muromachi Kikai, Japan) to measure the freezing of mice. All sessions were recorded using Bandicam software (Bandisoft, Korea) or AG-desktop recorder software (T. Ishii, Japan). The cumulative duration (s) spent in the complete absence of movement, except for respiration, was considered to be the freezing duration. Automated scoring of the freezing response was initiated after 1 s of persistent freezing behavior, and the freezing data of optogenetic and imaging experiments were manually reanalyzed by other non-behavioral operators (E.M. and R.O.S.) in blinded condition with the same criteria, to exclude the effect of optogenetic device (optic fiber for wired optogenetics or Teleopt battery for wireless optogenetics) and calcium imaging device attachments on automated animal tracking.

### Light fear conditioning (LFC) task

LFC was conducted under dim light (approximately 2 lx) conditions as described previously with some modifications^19^. Two distinct contexts were used for LFC habituation, training, and testing sessions. For habituation, training, and contextual fear test sessions, a triangle-type chamber (context A: Cxt-A) with black stripe patterns was used. This chamber was a triangular prism (one side × height: 180 × 250 mm), with a transparent acrylic board for the front wall, black stripe-patterned side walls with an 8 W white light bulb, and a floor made from 26 stainless steel rods. For cued-fear test sessions, a quadrangular prism chamber (width × depth × height: 190 × 180 × 420 mm, respectively) was used (context B: Cxt-B), with a transparent acrylic front board and white side walls with an 8 W white light bulb and an asperity white floor. Before each test session, the white floor was scented with 0.25% benzaldehyde water. For the habituation session, mice were allowed to explore the Ctx-A apparatus for 6 min per day for 2 days (non-operated, wired optogenetic, and imaging experiments) or for 3 days (wireless optogenetic experiment), and were then returned to their home cages.

For the training session 1 day after the habituation session, mice were conditioned in Cxt-A by ten pairings of the light-conditioned stimulus (CS) for 15 s with the unconditioned stimulus (US) (3 s footshock at the end of CS presentation, 0.4 mA) at 30 s intervals after a 150 s acclimation time. For the light-cued-fear memory test session, which was conducted 2 and 24 h after conditioning, different experimental mice were placed in Cxt-B for 120 s and then received two presentations of CS for 30 s at intervals of 30 s. For the contextual fear memory test session, 24 h after conditioning, different experimental mice were placed in Cxt-A for 360 s (non-operated animal experiments), while the same mice being tested for cued-fear memory were placed in Cxt-A for 360 s 1 hour after the light-cued-fear memory test (wireless optogenetic experiment).

### Auditory fear conditioning (AFC) task

AFC was conducted under normal light conditions as described previously with some modifications^43^. Two distinct contexts, described above, were used for AFC habituation, training, and testing sessions. For habituation and training sessions, a triangle-type chamber, Cxt-A, was used. This chamber had a transparent acrylic board for the front wall, black stripe-patterned side walls with a speaker, and a floor made from 26 stainless steel rods. For cued-fear test sessions, a quadrangular prism chamber, Cxt-B, was used. This chamber had a transparent acrylic front board and white side walls with a speaker and an asperity white floor.

For the 2 day habituation session, mice received four tone CS presentations (CS: 30 s at intervals of 30 s, 7 kHz, and 75 dB) after 120 s of exposure to Cxt-A, and then were returned to their home cages. For training sessions 1 day after the habituation session, mice were conditioned in Cxt-A by four pairings of the tone CS for 30 s with the US (1 s footshock at the end of CS presentation, 0.4 mA) at 30 s intervals after a 120 s acclimation time. For the tone-cued-fear memory test session, 24 h after conditioning, mice were placed in Cxt-B for 120 s and then subjected to four CS presentations for 30 s at intervals of 30 s.

### Pre-exposure facilitated contextual fear conditioning (pre-exposure facilitated CFC) task

Pre-exposure-facilitated CFC was conducted as described previously with some modifications^21^. A quadrangular prism chamber, Cxt-B, with a footshock grid as described above was used. On day 1, mice were placed in Cxt-B for 6 min and then were returned to their home cages. On day 2, mice were placed again in Cxt-B and immediately given the US (1 s footshock); kept for 10 s in the context; and then returned to their home cage. On day 3, to assess contextual freezing, mice were placed in Cxt-B again for 360 s.

### Optogenetic experiments

Wired and wireless optogenetic experiments were conducted as described previously with some modifications^29^. For the wired optogenetic experiment, on the recall session, mice were anesthetized with 3% isoflurane for placement of the optical fiber units, and dummy cannulae were removed from the guide cannulae. The black-stained optical fiber unit, comprising a plastic cannula body, was a two-branch-type unit with a black-stained optic fiber diameter of 0.250 mm (COME2-DF2-250; Lucir, Japan). The optical fiber unit was inserted into the guide cannulae, and the guide cannulae and the optical fiber unit were tightly connected with the optical fiber caps (303/OFC, Plastics One). The tip of the optical fiber was targeted slightly above the hippocampal CA3 (from bregma: -2.0 mm AP, ± 2.2 mm ML; from dura: + 1.2 mm DV). Mice attached with an optical fiber were then returned to their home cages and left individually at least for 1 h before beginning the cued-fear test session. Immediately before beginning the test session, mice were moved to the experimental room, and the fiber unit connected to the mouse was attached to an optical swivel (COME2-UFC, Lucir), which was connected to a laser (200 mW, 589 nm, COME-LY589/200; Lucir) via a main optical fiber. The delivery of light pulses was controlled by a custom-made schedule stimulator with OpenEx Software Suite (RX8-2, Tucker Davis Technologies, USA) in synchronized mode with a behavioral video tracking system (Muromachi Kikai). During both LFC and AFC test sessions, optical illumination (continuous 589 nm light, approximately 5 mW output from the fiber tip) was delivered to the CA3 concurrently with the onset of the first CS in both LFC or AFC tasks, and maintained until the end of the test sessions. Mice were then returned to their home cages individually, and then the attached optic fiber was removed from the mice after the anesthesia. For post-hoc analysis, mice were deeply anesthetized with a mixed anesthesia solution as described above, and perfused transcardially with 4% paraformaldehyde in PBS (pH 7.4), followed by immunohistochemical analysis to confirm virus vector infection.

For the wireless optogenetic experiment, to allow habituation to the 2 gram battery units (Bio Research Center), attachment and removal of the battery units was initiated from the habituation session by anesthetizing mice with 3% isoflurane before and after each behavioral session, respectively. The battery unit was attached to the implanted Teleopt LED device above the head, and mice were then returned to their home cages and left individually for at least 1 h until initiating behavioral sessions. For habituation to Cxt-A, mice attached to battery units were placed in Cxt-A for 10 min per day for 3 days. The time for pre-context habituation in the wireless optogenetic experiment was extended compared to the non-operated and wired optogenetic experiments to get mice attached with the battery well habituated to the chamber, because the battery’s width was a little bit bigger than the widths of parts of optogenetic guide cannula and mice head. After 3 days of habituation sessions, mice were subjected to ten CS-US pairings of LFC in the training session. During the LFC training session, wireless optical illumination (590 nm continuous light, approximately 10 mW output from the fiber tip) was delivered to the CA3 region ten times for 10 s at 35 s intervals, during either the inter-trial interval phase following CS-US presentation (Tr-ON group) or during resting in the home cage 10 min after the training session (HC-ON group). Mice were then returned to their home cages individually, and the battery unit was removed under anesthesia. The delivery of light pulses was controlled by a custom-made schedule stimulator system as described above. One day after the training session, mice were attached to the battery unit and then subjected to a cued-fear memory test followed by a contextual fear memory test at 1 h intervals. For post-hoc analysis, mice were deeply anesthetized with the mixed anesthesia solution described above and perfused transcardially with 4% paraformaldehyde in PBS (pH 7.4), followed by immunohistochemical analysis to confirm virus vector infection.

### *In vivo* Ca^2+^ imaging data acquisition in freely moving and head-fixed animals

Attachment and removal of a microendoscope was performed under 3% isoflurane anesthesia before and after each behavioral experiment. Mice attached to the microendoscope were returned to their home cages to recover for at least 30 min before and after the behavioral session. For the freely moving imaging experiment, mice were habituated to the endomicroscope attachment for 10 min per day for 3 days in their home cages before beginning the LFC behavioral study. During the 2 day habituation session, mice were also attached to the microendoscope. In both habituation sessions using the home cage and behavioral context, calcium imaging was performed, but acquired data were not analyzed. Subsequently, actual imaging began from the LFC training session to the test sessions. For freely moving CA1 Ca^2+^ imaging, mice were subjected to training, 30 min resting after training, the 2 h short-term memory (STM) test, and then the 24 h long-term memory (LTM) test using the same LFC protocol.

During the resting session, imaging was performed for 30 min. During the STM and LTM test sessions, after a 120 s acclimation session, mice were subjected to four CS presentations for 30 s at intervals of 30 s. However, STM session data were not analyzed because they were not important for the conclusion in this study. For freely moving CA3 Ca^2+^ imaging, mice were subjected to training, allowed to rest, and the 24 h LTM test was conducted using the same LFC protocol. During the resting session, imaging was performed for 30 min. During the LTM test sessions, after a 120 s acclimation session, mice were subjected to four CS presentations for 30 s at intervals of 30 s.

For the head-fixed imaging experiment, mice were habituated to endomicroscope attachment, and the fixation to the head-fixed apparatus comprised a footshock grid and an 8 W white light bulb, covered with a hemi-square paper box for 10 min per day for 4 days before beginning the head-fixed experiment. During habituation sessions, calcium imaging was performed, but acquired data were not analyzed. The footshock grid and light bulb were identical to the devices used for the freely moving LFC paradigm, and were measured with behavioral software (Muromachi Kikai). On the next day, mice were subjected to a CS (15 s constant light) presentation session and then a US (3 s footshock, 0.4 mA) presentation at intervals of 1 h. For the CS presentation session, after 150 s acclimation, mice were subjected to ten CSs for 15 s at 30 s intervals. For the US presentation session, after a 162 s acclimation time, mice were subjected to ten USs for 3 s at 42 s intervals. The durations of CS and US presentation sessions were 10 min total. In both freely moving and head-fixed imaging experiments, Ca^2+^ imaging was performed under dim light (approximately 2 lx) conditions, and the onset of behavioral and imaging systems was synchronized using the OpenEx Software Suite (RX8-2, Tucker Davis Technologies). Ca^2+^ signals produced from G-CaMP7 protein expressed in CA3 and CA1 excitatory neurons were captured at 20 Hz with nVista acquisition software (Inscopix) at the optimal gain and power of nVista LED. Ca^2+^ imaging movie recordings of all behavioral sessions were then extracted from the nVista Data acquisition (DAQ) box (Inscopix).

For post-hoc analysis, mice were deeply anesthetized with the mixed anesthesia solution described above and perfused transcardially with 4% paraformaldehyde in PBS (pH 7.4), followed by immunohistochemical analysis to confirm virus vector infection.

### *In vivo* Ca^2+^ imaging data processing and analysis

In both freely moving and head-fixed imaging experiments, only completely motion-corrected data was used for subsequent analysis. Data with an inadequate frame or that could not be corrected were excluded from analysis. Using Inscopix data processing software (IDPS, Inscopix) to create a full movie, recorded raw movies were temporally concatenated, spatially down-sampled (2×) and cropped, and then corrected for motion artifacts against a reference frame. A reference frame showing clear blood vessels as landmarks was chosen, and other frames were then aligned to the reference frame. Further motion correction was performed using Inscopix Mosaic software (Mosaic, Inscopix) as described previously^41, 44^. The corrected full movie was then temporally divided into individual behavioral sessions using Inscopix Mosaic software.

Subsequently, each individual session movie was low bandpass-filtered to reduce noise using Fiji software (NIH, USA) as described previously (see Extended Data Fig. 2). The change of fluorescence signal intensity (ΔF/F) for each behavioral session was subsequently calculated using Inscopix Mosaic software according to the formula ΔF/F = (F – Fm)/Fm, where F represents the fluorescence of each frame and Fm is the mean fluorescence for the entire session movie. Subsequently, movies representing each session were re-concatenated to generate full movies including all sessions in the ΔF/F format. Finally, cells were identified using an automatic sorting system, and HOTARU and Ca^2+^ signals of the detected cells over time were extracted in a (Ď; time × neuron) matrix format, as described previously.

Subsequent data processing and analysis were performed using a custom-made MATLAB code. Ca^2+^ signals were subjected to high-pass filtering (> 0.01 Hz threshold) to remove low frequency fluctuations and background noise in each Ca^2+^ cell signal, in which negative values were replaced with “0”. Using the filtered Ca^2+^ signal, the z-scores of behavioral sessions (training, resting, and LTM test sessions) were separately calculated to normalize and detect Ca^2+^ activities by thresholding (> 3 Standard Deviations from the ΔF/F signal) at the local maxima of the ΔF/F signal^44^. Then, to calculate the responsiveness of the cells to each behavioral event, z-scored Ca2^+^ activity was temporally sorted into nine behavioral events consisting of training acclimation (baseline), training CS (CS), training US (US), inter-trial interval 1 (ITI-1; ∼0–15 s after US-CS), ITI-2 (∼16–30 s after CS-US) sessions in the training session, resting (Rest) session, test acclimation (Test-Acc), CS (Test-CS), and ITI (Test-ITI) in LTM test sessions. The mean Ca^2+^ activities corresponding to behavioral events were calculated, and then divided by each cell baseline event to index responsiveness across behavioral events. Afterwards, CS-, US-, and Test-CS-responsive subpopulations with 2× greater responsiveness to stimuli than that of the baseline event were sorted. Ca^2+^ activities of these subpopulations were tracked across the LFC paradigm to calculate the mean Ca^2+^ activities of nine behavioral sessions and 1 s average Ca^2+^ activities for subsequent analyses, by which Ca^2+^ activities between genotypes were compared.

For functional connectivity analysis, to detect the Ca2^+^ event, the z-scores in CS-, US-, and Test-CS-responsive subpopulations as described above were binarized by thresholding (> 3 Standard Deviations from the ΔF/F signal) at the local maxima of the ΔF/F signal, and then were temporally down-sampled from 20 to 2 Hz data (500 ms binning). Subsequently, the functional connectivity, consisting of the number of synchronized activities among neurons of the two subpopulations in each 500 ms time window, was calculated and normalized as the functional connectivity in each behavioral session. The equation used for this analysis is below:

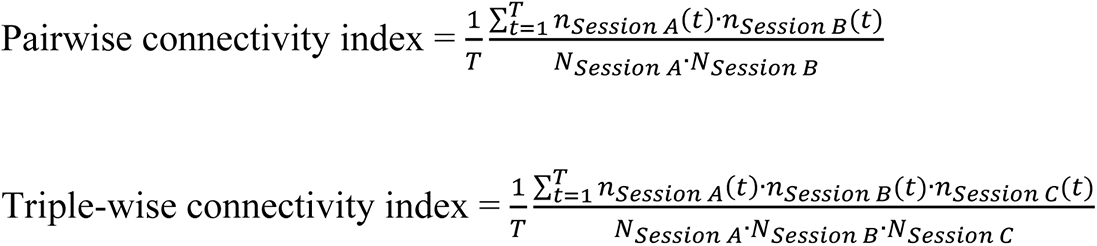

Where *n_Session A_*(*t*), (*n_Session B_*(*t*), (*n_Session C_*(*t*)) is the number of Session A (Session B, Session C) cells that were active in the time bin *t*; *N_Session A_* (*N_Session B_, N_Session_ C*) is the total number of Session A (Session B, Session C) cells; and is the total number of time bins for each session.

When the correlation between cued freezing and functional connectivity during ITI-1 was calculated, the variable, the change in ITI-1 connectivity relative to the baseline session in each cell, was calculated and used for correlation analyses.

### Population vector analyses

Calculations for the Mahalanobis population vector distance (PVD) and population vector rotation were conducted as described previously with some modifications. For Mahalanobis population vector distance, the 1 s-averaged z-scores in CS- and US-responsive subpopulations described above were used after principal component analysis (PCA)-based dimension reduction^26^. Since Mahalanobis distance does not work well because of the curse of dimensionality when the number of cells/dimensions (p) is greater than the number of available samples (n), (p > n), PCA was used to reduce the number of cells/dimensions in the data sets The data sets of 1 s-averaged z-scores in subpopulations are reduced into a lower dimension, and subsequently, the top three PCA scores (PC1, PC2, and PC3) are used to calculate the Mahalanobis population vector distance via PCA to quantify the similarity of two sets of neuronal representations between the CS or US session and ITI-1 sessions in CS and US ensembles, respectively. We defined a group of 3-dimensional activity vectors, , for each behavioral session (CS, US, or ITI-1) and calculated the PVD between the two representations. For example, the Mahalanobis PVD (*M*) between sets of CS- and ITI-1- evoked ensemble activity patterns in the CS ensemble is as follows:

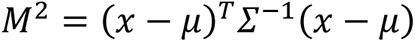

where *x* and *μ* are the individual and mean population vectors for the ITI-1 and CS ensemble activities, respectively, and *x^T^* and *μ^T^* are their transposes. The Mahalanobis distance accounts for differences in the means of the two sets of ensemble activities as well as their co-variances. The average PVD over all points in both sets of ensemble activities was calculated. To analyse the CS-ITI-1 and US-ITI-1 PVDs during the ten CS-US presentations in the CS and US ensembles, respectively, the top three PC scores calculated from the 1 s-averaged z-score data set are used, and the scores sorted by the CS, US, and ITI-1 sessions across ten trials are used for the mean population vector construction; subsequently, the relative change in PVD to ITI in each ensemble is calculated.

When the rotation of population vector between CS or US and ITI-1 sessions in CS- and US-responsive ensembles was calculated, we used the 1 s-averaged z-scores in CS- and US-responsive subpopulations.

### Immunohistochemistry and microscopy

Immunohistochemistry was conducted as described previously^29^. Mice were deeply anesthetized with the mixed anesthesia solution described above and perfused transcardially with 4% paraformaldehyde in PBS (pH 7.4). Brains were removed and further post-fixed by immersion in 4% PFA in PBS for 24 h at 4°C. Each brain was equilibrated in 25% sucrose in PBS for 2 days and then frozen in dry ice powder. Fifty μm coronal sections were cut on a cryostat and stored at -20°C in cryoprotectant solution (25% glycerol, 30% ethylene glycol, 45% PBS) until further use. For immunostaining, sections were transferred to 12-well cell culture plates (Corning, USA) containing Tris-buffered saline TBS-T buffer (with 0.2% Triton X-100, 0.05% Tween- 20).

For EYFP or G-CaMP and/or RGS-14 detection, after washing with TBS-T buffer, the floating sections were treated with blocking buffer (5% normal donkey serum [S30, Chemicon, USA] in TBS-T) at room temperature for 1 h. Primary antibody incubations were performed in blocking buffer containing rabbit anti-GFP (1:500, A11122; Molecular Probes, USA) and/or mouse anti-RGS-14 (1:500, N133/21; NeuroMab, USA) antibodies at 4°C for 1–2 days. After three 20 min washes with TBS-T, the sections were incubated with donkey anti-rabbit IgG-AlexaFluor 488 (1:500, A21206; Molecular Probes) and/or donkey anti-mouse IgG-AlexaFluor 546 secondary antibodies (1:500, A11036; Molecular Probes) in the blocking buffer at room temperature (RT) for 3 h.

For tdTomato detection, after washing with TBS-T buffer, floating sections were treated with blocking buffer (5% normal goat serum [S1000, Vector Laboratories, USA] in TBS-T) at RT for 1 h. Incubation with primary antibodies was performed in blocking buffer containing rabbit anti-DsRed (1:1000, 632496; Clonetech-Takara Bio, Japan) antibody at 4°C for 1–2 days. After three 20 min washes in TBS-T, sections were incubated with goat anti-rabbit IgG-AlexaFluor 546 secondary antibodies (1:300, A11035; Molecular Probes) in blocking buffer at RT for 3 h. Sections were treated with DAPI (1 μg/mL, 10236276001; Roche Diagnostics, Switzerland) and then washed with TBS three times (20 min/wash). The sections were mounted on slide glass with ProLong Gold antifade reagent (Invitrogen, USA). Images were acquired using a Keyence microscope (BIO-REVO, KEYENCE, Japan) with a Plan-Apochromat 4× or 20× objective lens.

### Statistics

Data are presented as means ± s.e.m. unless specified otherwise. Box plots represent median, first, and third quantiles, and their whiskers show minimum and maximum values. In box plots, outlier values are not shown for clarity of presentation, but all data points and animals were included in statistical analyses. Statistical analyses were performed using Excel (Microsoft) with Statcel4 (OMS, Japan) and MATLAB (Mathworks, USA) as described previously^29^. Comparisons of data between two groups were analyzed with a two-sided Student’s *t* test, Wilcoxon rank sum test, or Wilcoxon signed-rank test, based on the distribution and “n” size of the data. Correlation was analyzed with a Pearson correlation coefficient test. Multiple-group comparisons were conducted using two-way analysis of variance (ANOVA) with a post-hoc Tukey– Kramer multiple comparisons test when significant main effects were detected. Quantitative data are expressed as means ± SEM.

### Data and code availability

The data and codes that supported the findings of this study are available from the corresponding author upon reasonable request.

## Acknowledgments

We are grateful to K. Deisseroth (Stanford University) for providing eArch3.0-EYFP cDNA; S. Tonegawa (MIT) and S. Itohara (RIKEN) for providing floxed-NR1 transgenic mice; T. Fukai (Okinawa Institute of Science and Technology) and T. Haga (RIKEN) for mathematical analysis; T. Takekawa (Kogakuin University) for early access to the HOTARU detection system; M. Ito and N. Takino (Jichi Medical University) for production of the AAV vector. We thank alumni and current members of the Inokuchi laboratory for discussion. This work was supported by JSPS KAKENHI (grant number: JP18H05213), the Core Research for Evolutional Science and Technology (CREST) program (JPMJCR13W1) of the Japan Science and Technology Agency (JST), a Grant-in-Aid for Scientific Research on Innovative Areas “Memory dynamism” (JP25115002) from MEXT support to K.I., and the Grant-in-Aid for JSPS KAKENHI Scientific Research(B) (20H03554), Challenging Research (Exploratory) (17K19445), THE HOKURIKU BANK Grant-in-Aid for Young Scientists, the FIRSTBANK OF TOYAMA SCHOLARSHIP FOUNDATION RESERCH GRANT, the Takeda Science Foundation, the Tamura Science and Technology Foundation, and the Narishige Neuroscience Research Foundation support to M.N.

## Author contributions

M.N. and K.I. designed the experiments and wrote the manuscript. M.N. and E.M. performed the surgeries. M.N. performed behavioral experiments. M.N., E.M., and R.O.S. analyzed behavioral data. M.N. and S.O. performed calcium data analysis. M.N. wrote MATLAB codes. S.M. prepared the AAV. K.I. supervised the entire project.

---

**Extended Data Figure 1.**
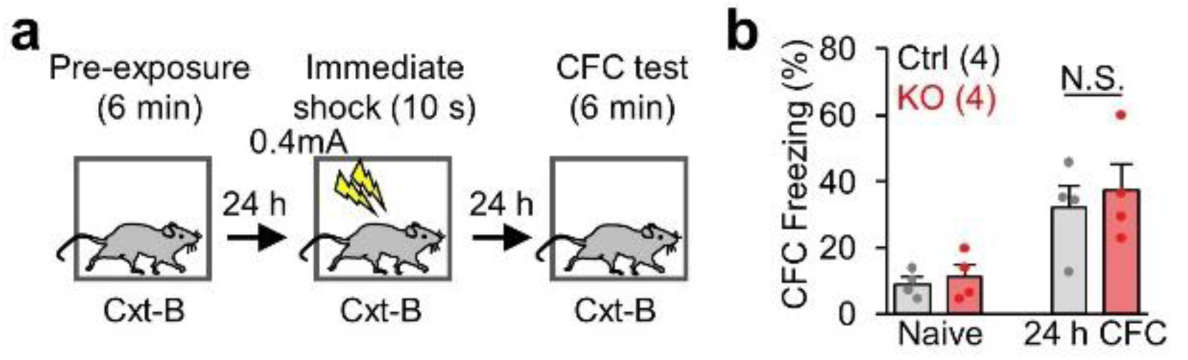
CA3-NR KO mice exhibited comparable contextual freezing in the pre-exposure-facilitated CFC task. **a,** Experimental design. **b,** Contextual freezing levels in 24 h long-term memory test. *P* values were calculated using an unpaired two-tailed *t* test. N.S., not significant (*P* > 0.05). Graphs represent means ± SEM, and circles in the graph represent individual animals. Numbers in parentheses denote the number of mice in each group used for the study. Lightning bolt, footshock; Cxt, context; HPC, hippocampus; AP, anterior-posterior; N.S., not significant.

**Extended Data Figure 2.**
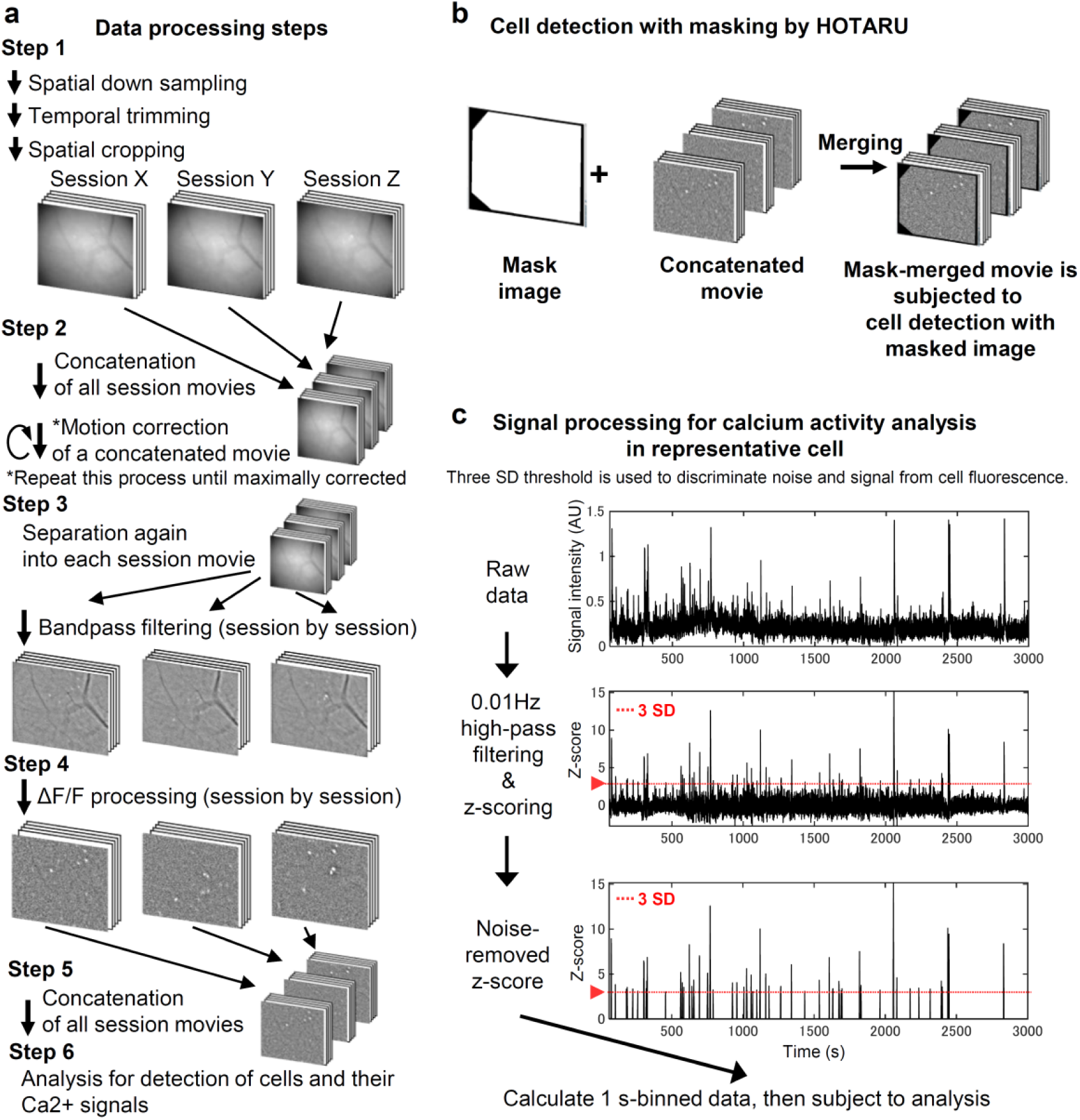
Data processing and cell detection for analysis. a, Data processing steps. Step 1: movies acquired from each behavioral session are down- sampled, temporally trimmed, and spatially cropped. Step 2: the movies are concatenated into a series of movies, and repeatedly corrected until artifacts of movement are minimized. Step 3: the concatenated movie is separated again into pre-concatenated movies and subsequently subjected to bandpass filtering. Step 4: dF/F conversion. Step 5: re-concatenation. Step 6: the concatenated- and registrated-movie is subjected to cell detection. **b,** A train of dF/F is subjected into the HOTARU algorithm to automatically detect active cells. **C,** The acquired calcium trace signal is converted into calcium activity by high-pass filtering, z-score normalization, and cutoff of inadequate signal in each cell to remove background fluctuation. Then 1 s-mean calcium activities are subjected to quantitative analyses. The dashed line indicates a cutoff threshold less than three standard deviations (< 3 SD).

**Extended Data Figure 3.**
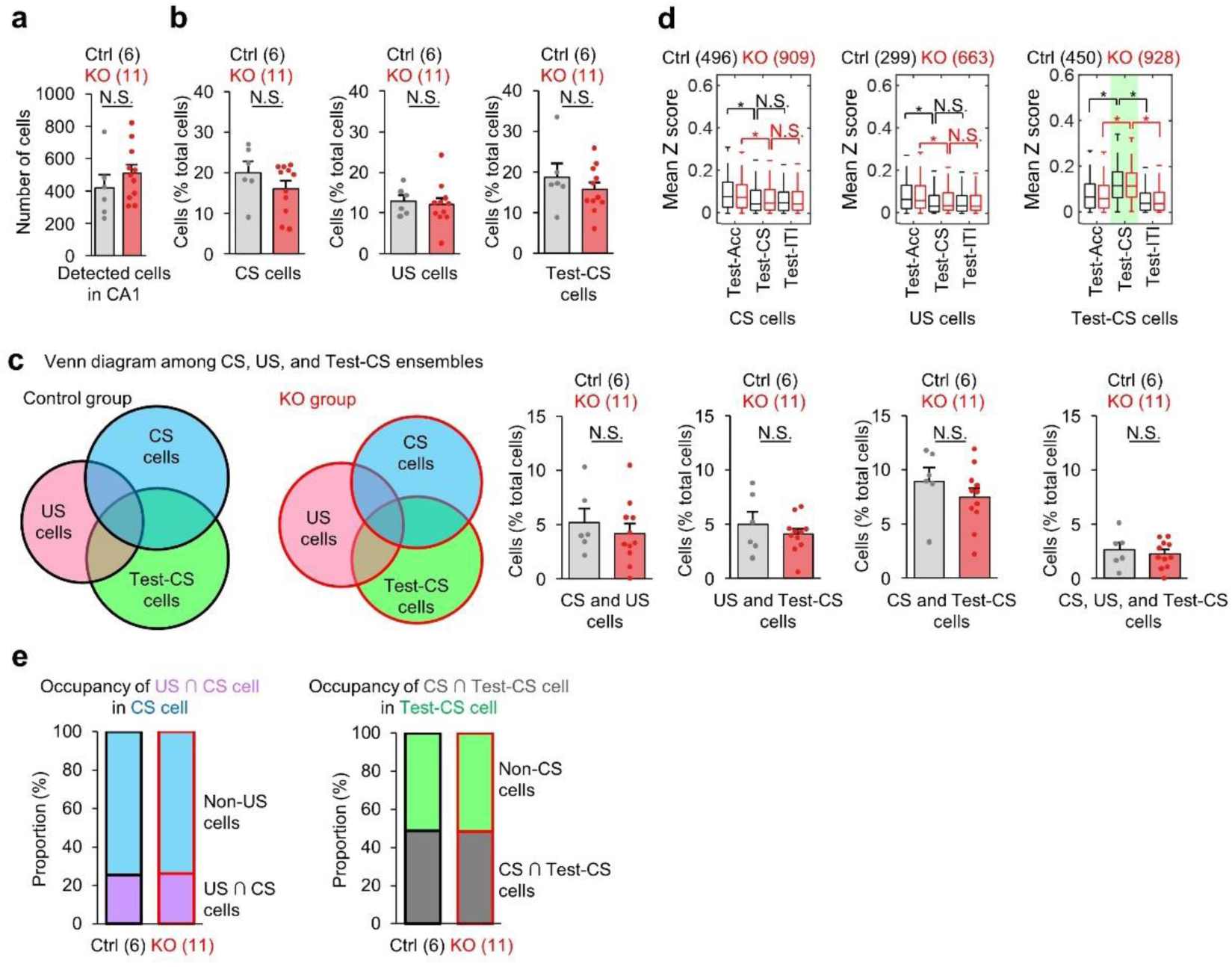
CA3-NR KO mice exhibited normal CA1 ensemble structure. a, Columns comparing number of detected cells during CA1 imaging in control and CA3-NR1 KO mice. **b,** Columns comparing percentile of ensemble size in CS-, US-, and Test-CS-responsive subpopulations between control and KO mice. **c,** Venn diagrams comparing and illustrating the overlapping and size of each ensemble in CA1. Columns comparing the percentiles of overlapping ensemble sizes among CS-, US-, and Test-CS-responsive subpopulations between control and KO mice. **d,** Box plots comparing mean z-scores of long-term memory test sessions between genotypes in CS-, US-, and Test-CS-responsive subpopulations. **e,** Left, occupancy of US ∩ CS-responsive cells in the CS-responsive cell population. Right, occupancy of CS-responsive cells in the Test-CS-responsive population. Numbers in parentheses denote the number of mice (**a-c**) or cells (**d**) in each group used for the study. *P* values were calculated using an unpaired two-tailed *t* test (**a-c, e**) or Wilcoxon signed-rank test (**d**) (\**P* < 0.001). N.S., not significant (*P* > 0.05). Box plots illustrate median, first, and third quantiles, and minimum and maximum values. Graphs represent means ± SEM, and circles in the graphs represent individual animals.

**Extended Data Figure 4.**
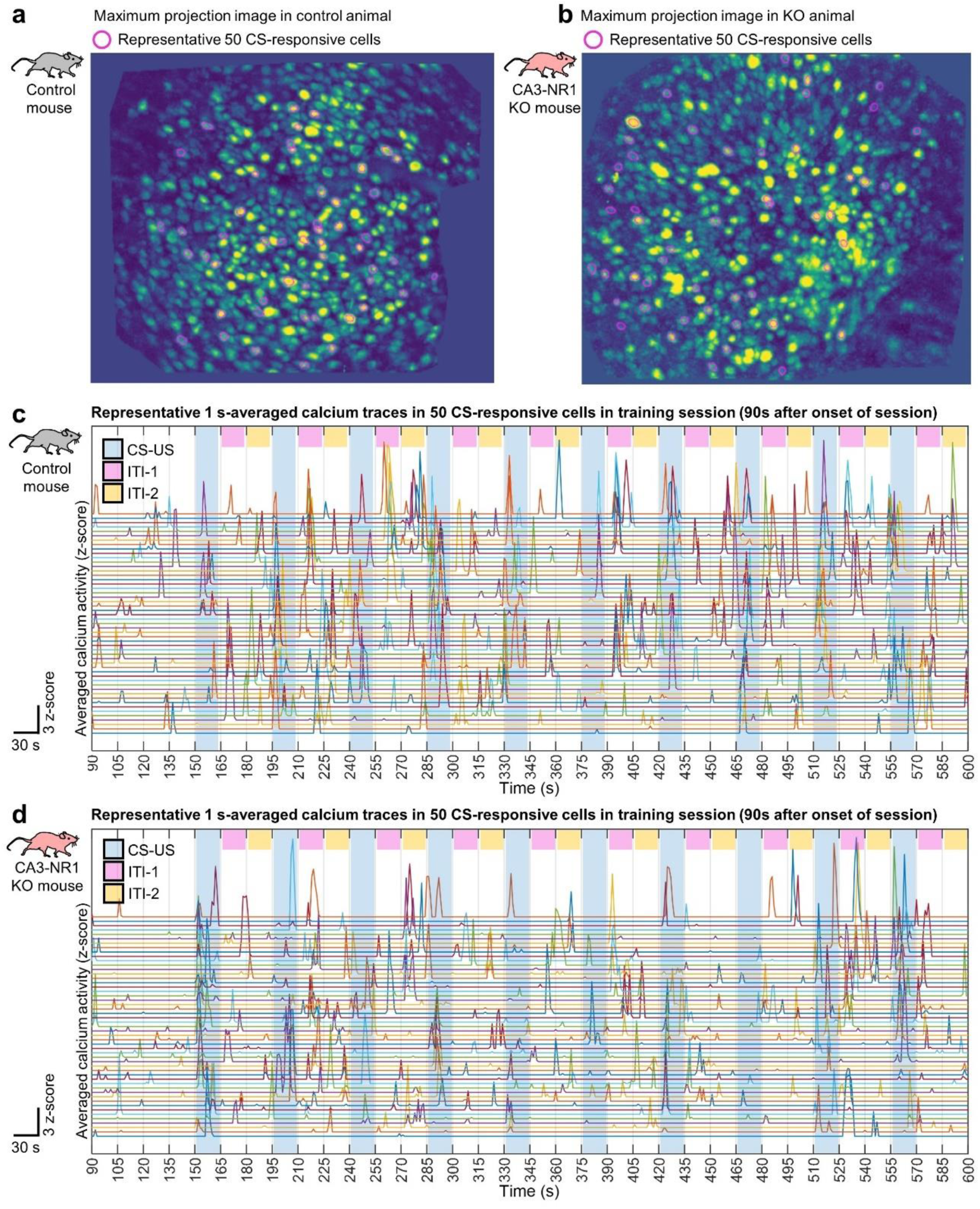
Representative *in vivo* calcium imaging data acquisition in CA1 of freely moving animals. **a, b,** Stacked- and pseudo-colored dF/F images acquired using the microendoscope over entire recording sessions of imaging in the hippocampus from control (**a**) and KO (**b**) animals. Magenta circles indicate the footprint contours of detected cells. **c, d,** Representative 1 s-averaged calcium activities in representative 50 CS-responsive cells during LFC training 90 s after onset of behavioral session to end in control (**c**) and KO (**d**) animals. Blue, pink, and yellow rectangles indicate the timings of CS-US, ITI-1, and ITI-2, respectively.

**Extended Data Figure 5.**
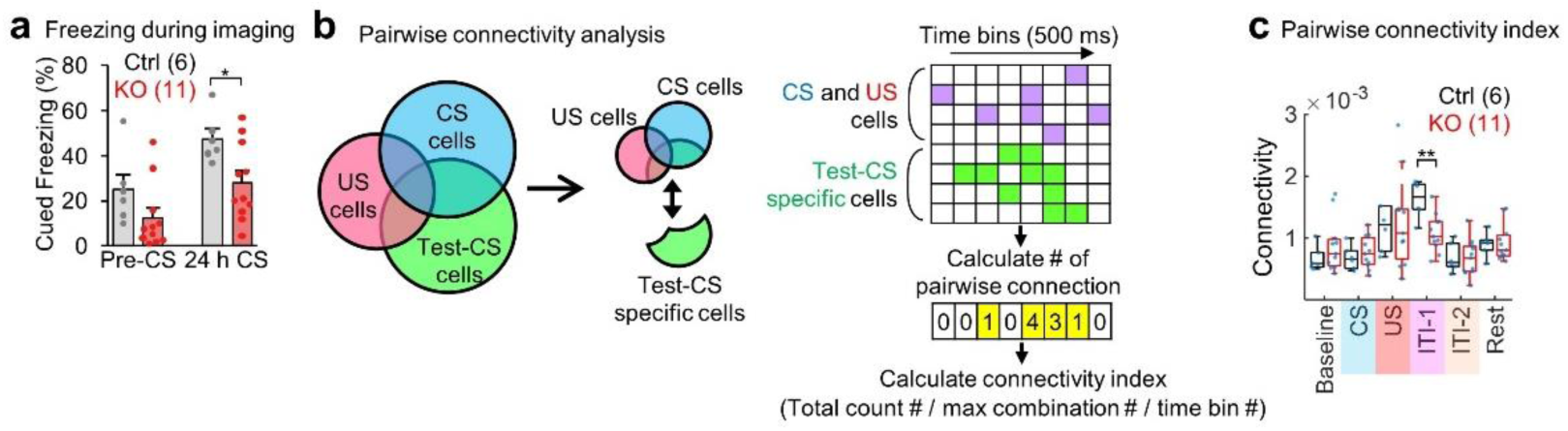
CA3-NR KO mice exhibited impaired functional connectivity between CS ∪ US- and Test-CS-responsive specific cells. **a,** Cued freezing levels during 24 h long-term memory tests in the imaging study. **b,** Scheme for connectivity analysis. Binarized Ca2^+^ activity in each cell is sorted into CS ∪ US- and Test-CS-responsive specific subpopulations, and then pairwise connectivity is calculated by normalizing the number of synchronized connections every 500 ms in each session. **c,** Box plots comparing the mean connectivity between genotypes in each session. Numbers in parentheses denote the number of mice (**a, c**) in each group used for the study. *P* values were calculated using an unpaired two-tailed *t* test (**a, c**) (\**P* < 0.05, \*\**P* < 0.01). Box plots illustrate median, first, and third quantiles, and minimum and maximum values. Graphs and scatter plots represent means ± SEM. In graphs, circles represent individual animals.

**Extended Data Figure 6.**
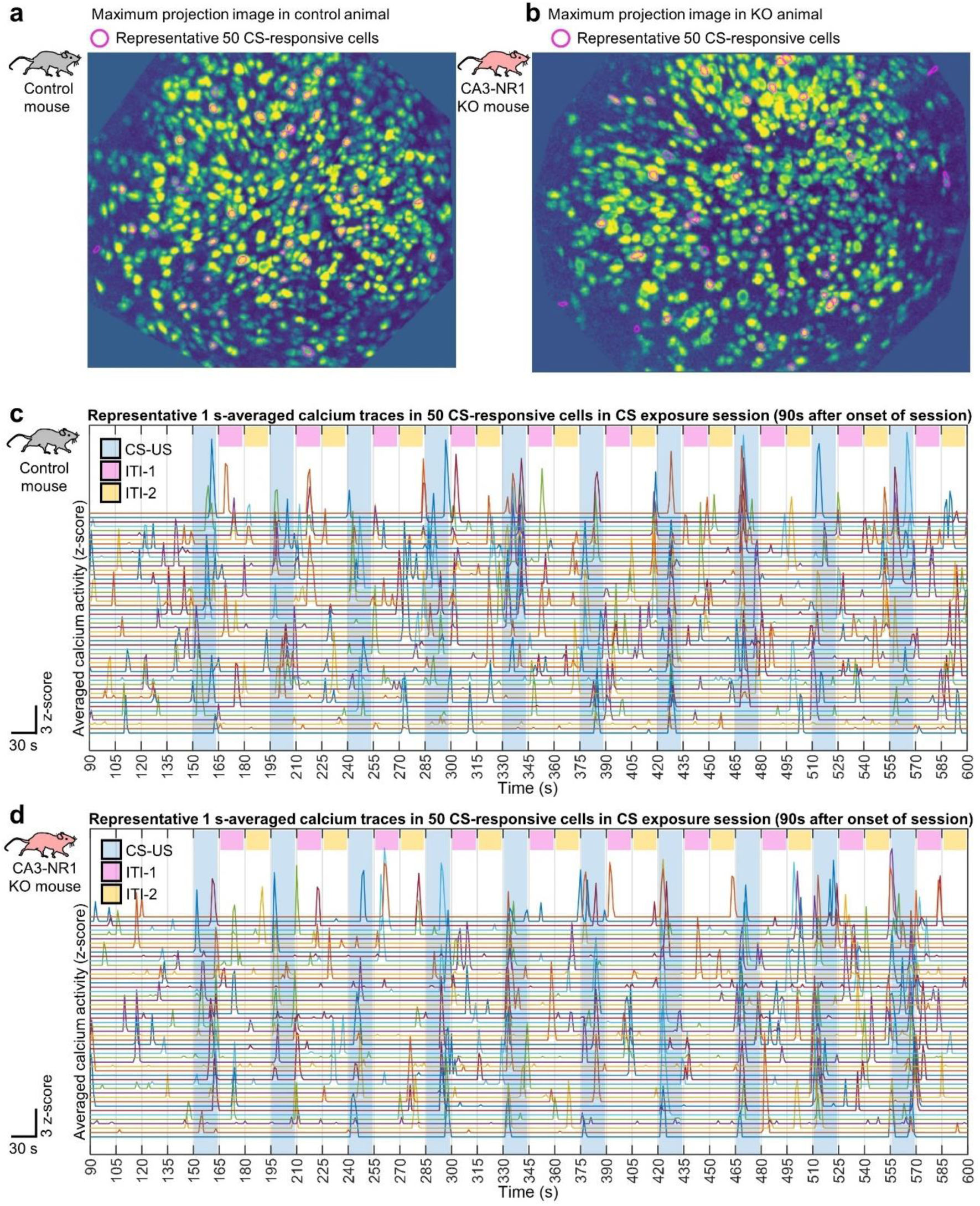
Representative *in vivo* calcium imaging data acquisition in CA1 of head-fixed animals. **a, b,** Stacked- and pseudo-colored dF/F images acquired through the microendoscope over entire recording sessions of imaging in the hippocampus (right) from (**a**) control and (**b**) KO animals. Magenta circles indicate the footprint contours of detected cells. **c, d,** Representative 1 s-averaged calcium activities in 50 representative CS-responsive cells during LFC training (90 s after beginning the behavioral session to the end) in (**c**) control and (**d**) KO animals. Blue, pink, and yellow rectangles indicate the timings of CS-US, ITI-1, and ITI-2, respectively.

**Extended Data Figure 7.**
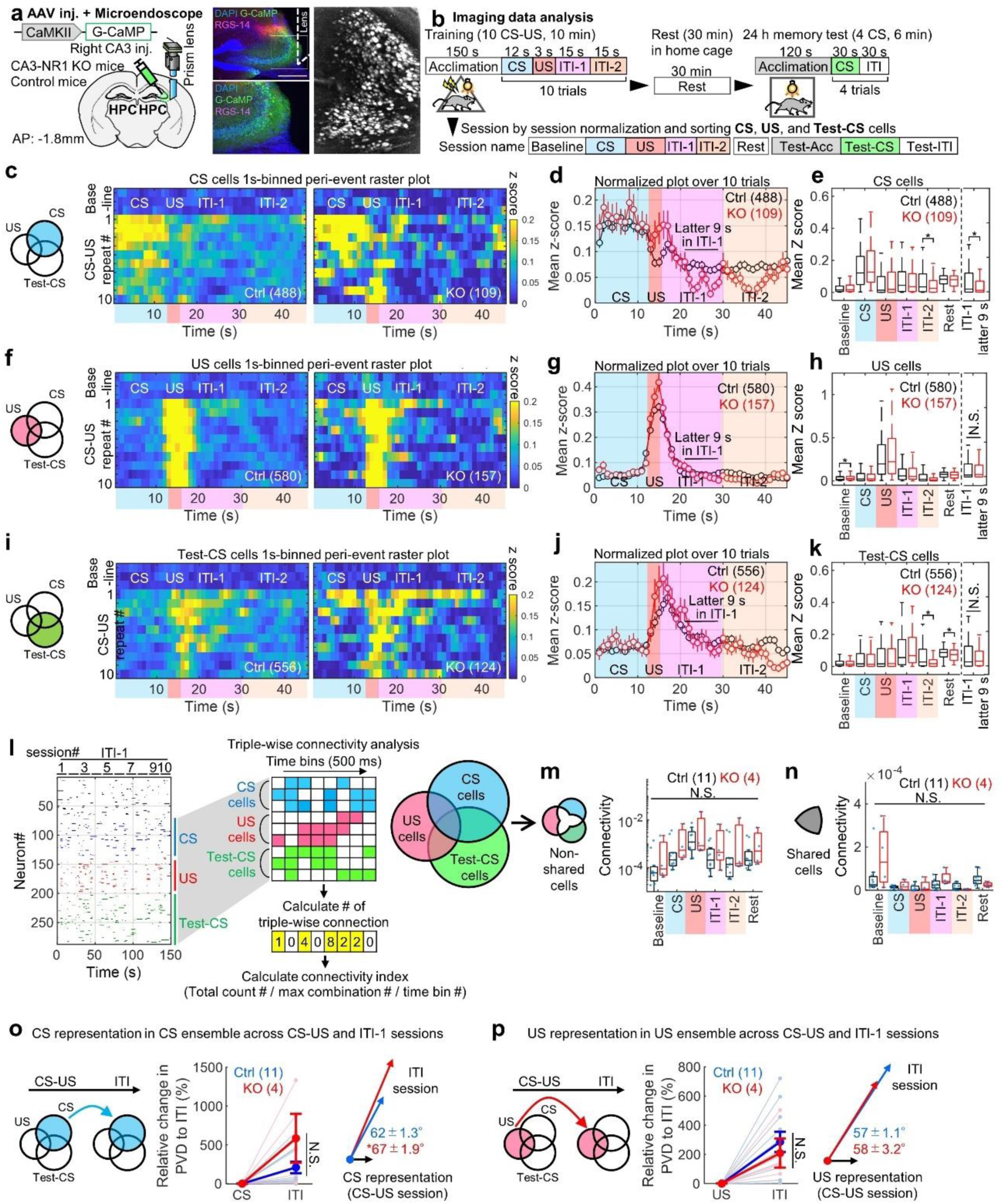
CA3-NR KO mice exhibited impaired reverberatory activity following CS-US presentation in CS- and Test-CS-responsive CA3 subpopulations under free-moving conditions. **a,** Left, experimental design. Right, coronal section of the hippocampus with G-CaMP-expressed cells (green) in CA3 and immunostained with anti-RGS-14 (red). RGS-14 is a marker for CA2 and stacked dF/F images acquired through the microendoscope over entire recording sessions of imaging in the hippocampus. Scale bar, 500 μm. **b,** Scheme of imaging data analysis. In each cell, Ca^2+^ data is classified into nine sessions, and the calculated mean z-score is considered to represent responsiveness and sorted into CS-, US-, and Test-CS-responsive subpopulations. **c,** Venn diagrams illustrating CS ensembles. Peri-event raster plots during training sessions in CS-responsive subpopulations of (left) control and (right) KO mice. Each short vertical tick represents a 1 s change of mean z-score across baseline and ten CS-US pairings. Ca^2+^ activities are aligned at the time when CS-US stimuli were delivered. The color code indicates mean z-scores. **d,** Averaged z-score plots over ten CS-US pairings in CS-responsive subpopulations. **e,** Box plots comparing mean z-scores between genotypes in each session. **f,** Venn diagrams illustrating ensembles. Peri-event raster plots during training sessions in US-responsive subpopulations of (left) control and (right) KO mice. Each short vertical tick represents a 1 s change of mean z-score across baseline and ten CS-US pairings. Ca^2+^ activities are aligned at the time when CS-US stimuli were delivered. The color code indicates mean z-scores. **g,** Averaged z-score plots over ten CS-US pairings in US-responsive subpopulations. **h,** Box plots comparing mean z-scores between genotypes in each session. **i,** Venn diagrams illustrating Test-CS ensembles. Peri-event raster plots during training sessions in Test-CS-responsive subpopulations of (left) control and (right) KO mice. Each short vertical tick represents a 1 s change of mean z-score across baseline and ten CS-US pairings. Ca^2+^ activities are aligned at the time when CS-US stimuli were delivered. The color code indicates mean z-scores. **j,** Averaged z-score plots over ten CS-US pairings in Test-CS-responsive subpopulations. **k,** Box plots comparing mean z-scores between genotypes in each session. **l,** Left, representative binarized raster plots of Ca^2+^ activity across ten ITI-1 sessions in control animals. Right, magnified raster plots focusing on CS-, US-, and Test-CS-responsive subpopulations and scheme for connectivity analysis. This analysis calculates connectivity by normalizing the number of synchronized connections in every 500 ms among three subpopulations in each session. **m, n,** Box plots comparing mean connectivity between genotypes in each session. **o, p,** Mahalanobis PVD and rotation between CS and ITI-1 sessions in the CS-responsive ensemble (**o**), and between US and ITI-1 sessions in the US-responsive ensemble (**p**). Numbers in parentheses denote the (**d, e, g, h, j, k**) number of cells or (**m, n, o, p**) mice in each group used for the study. *P* values were determined using a Wilcoxon rank sum test (**e, h, k**) or an unpaired two-tailed *t* test (**m, n, o, p**) (\**P* < 0.05). N.S., not significant (*P* > 0.05). Box plots indicate median, first, and third quantiles, and minimum and maximum values. Graphs indicate means ± SEM. In graphs, circles represent individual animals.

**Extended Data Figure 8.**
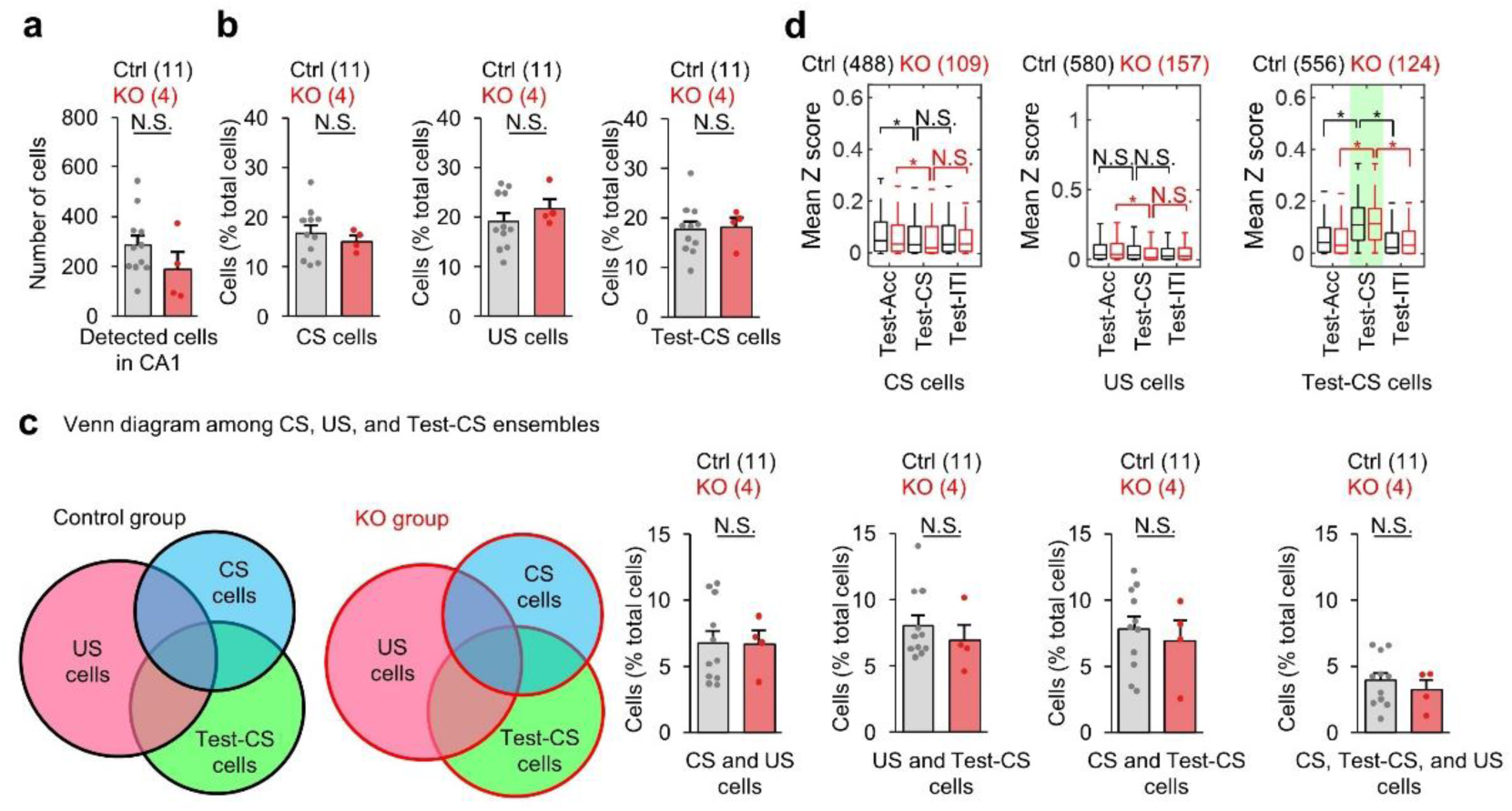
CA3-NR KO mice exhibited normalCA3 ensemble structure. **a,** Columns comparing the number of detected cells during CA3 imaging in control and CA3-NR1 KO mice. **B, C**olumns comparing percentiles of ensemble sizes in CS-, US-, and Test-CS-responsive subpopulations between control and KO mice. **c,** Venn diagrams comparing and illustrating the overlapping and size of each ensemble in CA3. Columns compare the percentiles of overlapping ensemble sizes between CS-, US-, and Test-CS-responsive subpopulations between control and KO mice. **D,** Box plots comparing mean z-scores of long-term memory test sessions between genotypes in CS-, US-, and Test-CS-responsive subpopulations. Numbers in parentheses denote the number of (**a-c**) mice or (**d**) cells in each group used for the study. *P* values were determined using (**a-c**) an unpaired two-tailed *t* test or (**d**) a Wilcoxon signed-rank test (\**P* < 0.001). N.S., not significant (*P* > 0.05). Box plots represent median, first, and third quantiles, and minimum and maximum values. Graphs show means ± SEM. In graphs, circles represent individual animals.

**Extended Data Figure 9.**
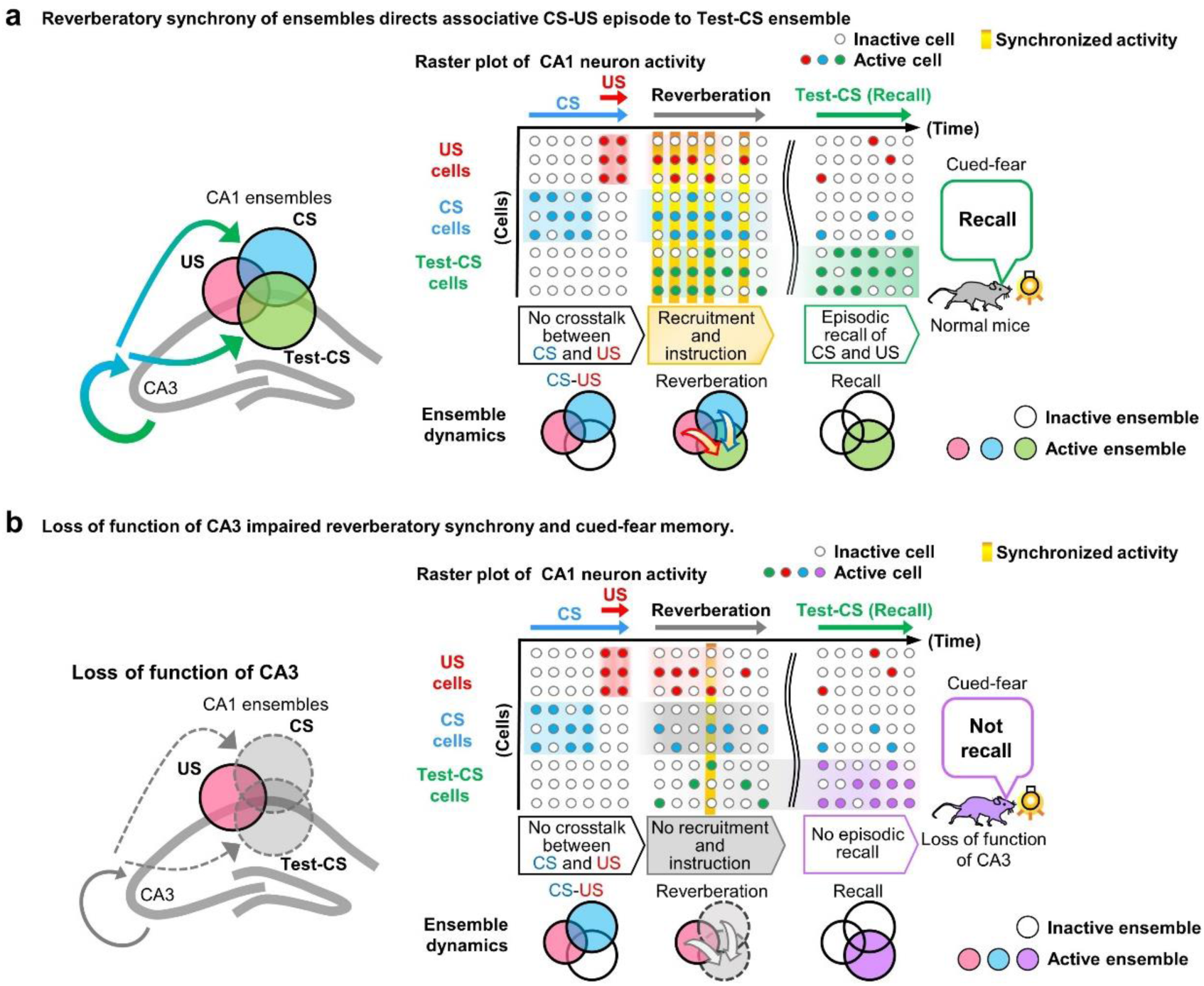
Model for cued-fear memory encoding in the hippocampal network using CA3-dependent reverberatory activity. **a,** left, Venn diagrams showing CS-, US-, and Test-CS-responsive cell ensembles in CA1 of normal mice, in which CA3-dependent reverberation occurs normally. Right, raster plots of CA1 subpopulations and with the timeline of the cued-fear memory paradigm. During CS and US inputs during training, CS and US information are separately encoded in CS- and US-responsive cell populations, respectively. During reverberation in training, co-activity of CS- and US-responsive cells recruits and instructs Test-CS-responsive cells in the CS-US episode. During recall, Test-CS-responsive cells drive the episodic recall of cued-fear memory. **b,** left, Venn diagrams of CS-, US-, and Test-CS-responsive ensembles in loss-of-function (CA3 NR KO and CA3 silencing during ITI). Right, during CS and US input in training, CS and US information are separately and normally encoded. However, without reverberation in training, the low co-activity frequency of CS- and US-responsive cells fails to recruit and instruct Test-CS-responsive cells in the episodic relation between CS and US. Thus, during recall, Test-CS-responsive cells fail to drive cued-fear memory recall. Filled circles with color indicate activated cells in each behavioral session. Arrows indicate the direction of information flow. Light bulb, light CS; yellow bar, moment-occurring synchrony among CS-, US-, and Test-CS cell ensembles.

**Extended Data Figure 10.**
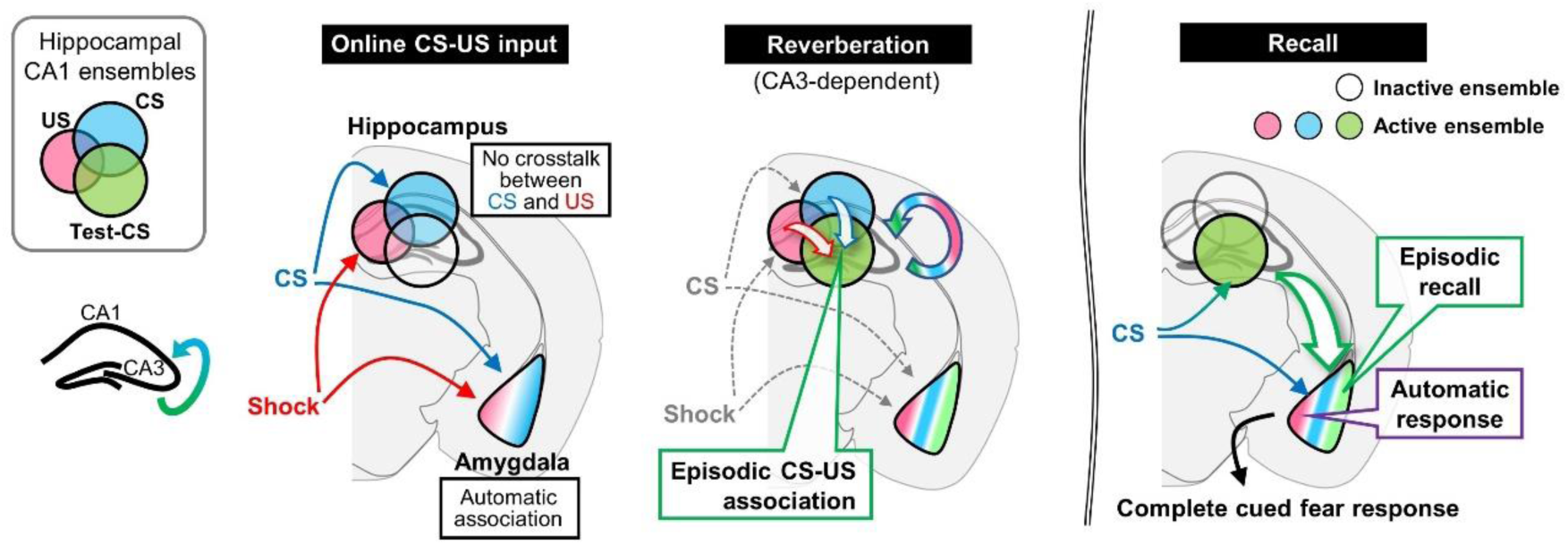
Model for hippocampal function in cued-fear memory. Venn diagrams showing CS-, US-, and Test-CS-responsive cell ensembles in CA1. During online CS and US inputs in training, the hippocampus encodes CS and US information independently, while the amygdala associates CS and US directly as automatic association. During reverberation, the hippocampus produces episodic CS-US association. During recall, the Test-CS-responsive cell ensemble in the hippocampus sends the episodic portion of the CS-US information to the amygdala to complete cued-fear memory. Filled circles and amygdala icons with color indicate temporal activation throughout learning and recall.

